# Biotic and abiotic drivers of ecosystem functioning differ between a temperate and a tropical region

**DOI:** 10.1101/2024.02.28.582312

**Authors:** Laura J. A. van Dijk, Andreia Miraldo, Dimby Raharinjanahary, Eric Tsiriniaina Rajoelison, Brian L. Fisher, Robert M. Goodsell, Elzbieta Iwaszkiewicz-Eggebrecht, David Åhlén, Johanna Högvall, Erika Lundberg, Emma Rova, Piotr Łukasik, Fredrik Ronquist, Tomas Roslin, Ayco J. M. Tack

**Author notes:** These authors share last authorship (Tomas Roslin, Ayco J. M. Tack).

## Abstract

Any single ecosystem will provide many ecosystem functions. Whether these functions tend to increase in concert or trade off against each other is a question of much current interest. Equally topical are the drivers behind ecosystem function rates. Yet, we lack large-scale systematic studies that investigate how abiotic factors can directly or indirectly — via effects on biodiversity — drive ecosystem functioning. In this study, we assessed the impact of climate, landscape and biotic community on ecosystem functioning and multifunctioning in the temperate and tropical zone, and investigated potential trade-offs among ecosystem functions in both zones. To achieve this, we measured a diverse set of insect-related ecosystem functions — including herbivory, seed dispersal, predation, decomposition and pollination — at 50 sites across Madagascar and 171 sites across Sweden, and characterized the insect community at each site using Malaise traps. We used structural equations models to infer causality of the effects of climate, landscape, and biodiversity on ecosystem functioning. For the temperate zone, we found that abiotic factors were more important than biotic factors in driving ecosystem functioning, while in the tropical zone, effects of biotic drivers were most pronounced. In terms of trade-offs among functions, in the temperate zone, only seed dispersal and predation were positively correlated, while all other functions were uncorrelated. By contrast, in the tropical zone, most ecosystem functions increased in concert, highlighting that tropical ecosystems can simultaneously provide a diverse set of functions. These correlated functions in Madagascar could for the most part be explained by similar responses to local climate, landscape, and biota. Our study suggests that the functioning of temperate and tropical ecosystems differs fundamentally in patterns and drivers. Without a better understanding of these differences, it will be impossible to correctly predict shifts in ecosystem functioning in response to environmental disturbances. To identify global patterns and drivers of ecosystem functioning, we will next need replicate sampling across biomes – as here achieved for two regions, thus paving the road and setting the baseline expectations.

## Introduction

The functioning of ecosystems relies on a wide variety of species and their interactions (Loreau et al. 2002, Tilman et al. 2014). For example, many plants are dependent on their pollinators for fertilization, and microbes and invertebrates in soil and litter are vital for nutrient cycling via the decomposition of organic matter (Ollerton et al. 2011). While it is well established that climate can affect biodiversity, our understanding of how abiotic and biotic conditions can jointly affect ecosystem functioning remains superficial (Bellard et al. 2012, Hooper et al. 2012). Specifically, studies simultaneously exploring the effects of climate, landscape, and biodiversity on various ecosystem functions are scarce, and often conducted at small spatial scales or under experimental conditions (van der Plas 2019). To identify generalizable links between climate, landscape, biodiversity and ecosystem functioning, we need extensive studies in natural settings. Such studies should monitor biodiversity and ecosystem functioning on large geographical scales and across climatic zones. Given that human-induced changes in climate, landscape and biodiversity have become increasingly noticeable across the globe (Walther et al. 2002), understanding their potential large-scale impacts on ecosystem functioning is urgently needed to safeguard healthy ecosystems.

Of particular interest is the relative impact of abiotic versus biotic impacts on ecological functioning in different biomes. Key theory (MacArthur 1984) and empirical observations (Pennings and Silliman 2005, Roslin et al. 2017, Baskett and Schemske 2018) suggest that biotic interactions are generally stronger in tropical regions than temperate regions, thus accentuating the role of biotic impacts at lower latitudes. The greater availability of energy near the Equator contributes to these patterns, and increases the activity of ectotherms such as arthropods (Roslin et al. 2017). Ecosystem function rates at lower latitudes may also be expected to increase, since higher temperatures promote chemical reaction speeds and enzymatic activities, which accelerate microbial functions like decomposition (Rubenstein et al. 2017). Yet, while the effects of latitude and temperature have been explored for individual ecological functions (Pennings and Silliman 2005, Roslin et al. 2017, Baskett and Schemske 2018), their role in determining ecosystem multifunctionality remains to be established.

Beyond latitudinal imprints, ecosystem functioning can be affected by characteristics of the landscape (Lovett et al. 2005, Gaitán et al. 2014). Vegetation cover is important, as it buffers the microclimate from variation in the regional climate, e.g. by reducing temperature and promoting humidity, with potential consequences for ecosystem functioning (Hardwick et al. 2015). Vegetation cover can also indirectly affect functioning by altering the diversity and composition of biotic communities (Peters et al. 2019). Notably, the effect of landscape may vary across climatic zones: In colder regions, blocking of sunlight by vegetation may reduce activity rates of ectotherms, leading to lower functional rates, while in warmer regions, blocking of sunlight may instead protect biota against desiccation and extreme heat, thus promoting functional rates. To pinpoint the effects of climate, landscape and biota on ecosystem functioning across zones, we need large-scale yet spatially well-replicated experiments that investigate ecosystem functioning within and across multiple climatic zones. Moreover, to disentangle direct and indirect effects of abiotic and biotic factors, we need to combine such large-scale experiments with statistical approaches that can infer the causality of drivers of ecosystem functioning.

While a positive relationship between biodiversity and ecosystem functioning has been confirmed in numerous experimental studies (Tilman et al. 2014), such studies are much scarcer in natural settings (Maestre et al. 2012, van der Plas 2019). Experimental studies exploring the link between biodiversity and ecosystem functioning are typically done under highly controlled environmental conditions which simplify the natural complexity of ecosystems (Hooper et al. 2005). In a natural context, relationships between biodiversity and ecosystem functions are more variable than experimental settings, and neutral relationships often outweigh positive ones (van der Plas 2019). For example, studies conducted in a natural setting show conflicting results regarding the effects of species richness on functioning, where species richness is reported to either decrease (Dee et al. 2023), increase (Duffy et al. 2017) or have no effect (Dormann et al. 2019) on plant productivity. To expand our understanding from experimental studies to the natural context, we need to accumulate studies on ecosystem functioning in natural settings, which are conducted over large spatial scales to discern the differences in drivers between biomes.

Since different ecosystem functions can respond differently to abiotic or biotic drivers, a substantial body of recent literature has aimed to uncover the drivers contributing to the *overall* functionality of an ecosystem, i.e., ecosystem multifunctionality. Abiotic as well as biotic drivers are expected to contribute to the multifunctionality of an ecosystem. For example, a highly biodiverse ecosystem is likely to simultaneously support multiple functions, as more taxa can potentially fulfil more functions (Perkins et al. 2015). Moreover, as temperature is generally expected to positively affect functional rates, higher rates of functions are more likely to occur at warmer sites, thus promoting the overall functionality of the ecosystem (Pan et al. 2017).

The multifunctionality of an ecosystem can potentially decrease when functions trade off against one another. For example, an ecosystem with high herbivory rates may experience lower pollination rates, since leaf loss and defence induction can reduce resources allocated towards the production of nectar and pollen (Jacobsen and Raguso 2018). On the other hand, functions could also increase in concert, potentially promoting ecosystem multifunctionality. For example, an ecosystem with low herbivory rates could experience lower decomposition rates, since unpalatable leaves also tend to decompose more slowly (Moretto et al. 2001). Besides direct mechanistic links among functions, positive or negative correlations among functions in space can potentially be driven by their synchronous or asynchronous responses to abiotic or biotic conditions. In colder regions like the temperate zone, many functions may be positively affected by warmer local temperatures, potentially leading to positive correlations among functions (Fig. S1). On the other hand, in warmer regions like the tropics, local temperature may have little influence on functionality, as the high regional temperature sustains high functional rates across all sites (Fig. S1). Yet, we do not know how such patterns vary across latitudinal gradients, as studies on ecosystem multifunctionality are often conducted at a local (Grime et al. 1996) or regional scale (Allan et al. 2015, Soliveres et al. 2016), while few studies of drylands cover multiple climatic zones (Maestre et al. 2012, Delgado-Baquerizo et al. 2016). Hence, we lack a systematic analysis of ecosystem multifunctionality and functional trade-offs in tropical and temperate zones, and we lack insights into potential differences in abiotic and biotic drivers acting within these zones.

In this study, we experimentally assessed and compared the climatic, landscape and biotic drivers behind ecosystem functioning and multifunctioning, and estimated positive and negative correlations among functions. We assessed ecosystem functioning in two regions from different climate zones – one temperate and one tropical region – across multiple natural sites. Specifically, we asked the following questions:

- What are the abiotic and biotic drivers behind ecosystem functioning and multifunctioning in a temperate and a tropical climate, and do they differ in strength between biomes?
- Do ecosystem functions increase in concert, or trade off against each other? In other words, are there positive or negative correlations among functions? Can these correlations be explained by responses to abiotic or biotic factors?

To this end, we measured a set of ecosystem functions at 72 to 171 sites across Sweden and 50 sites across Madagascar, including experiments on herbivory, seed dispersal, predation, decomposition and pollination, and characterized the insect community at each site using Malaise traps. As we aimed to describe the causality of drivers of ecosystem functioning, thereby disentangling both direct and indirect drivers, we adopted the approach of structural equation modelling (Lefcheck 2016).

## Materials and methods

### Study area

To resolve the drivers behind ecosystem functioning across climatic zones, we took measurements across the full latitudinal and longitudinal range of a temperate country, Sweden, as well as a tropical country, Madagascar (Fig. 1). At each of the sites, we conducted experiments to estimate ecosystem functions, characterized the insect community using Malaise traps, and recorded several climate and landscape variables, as further detailed below.

**Figure 1.**
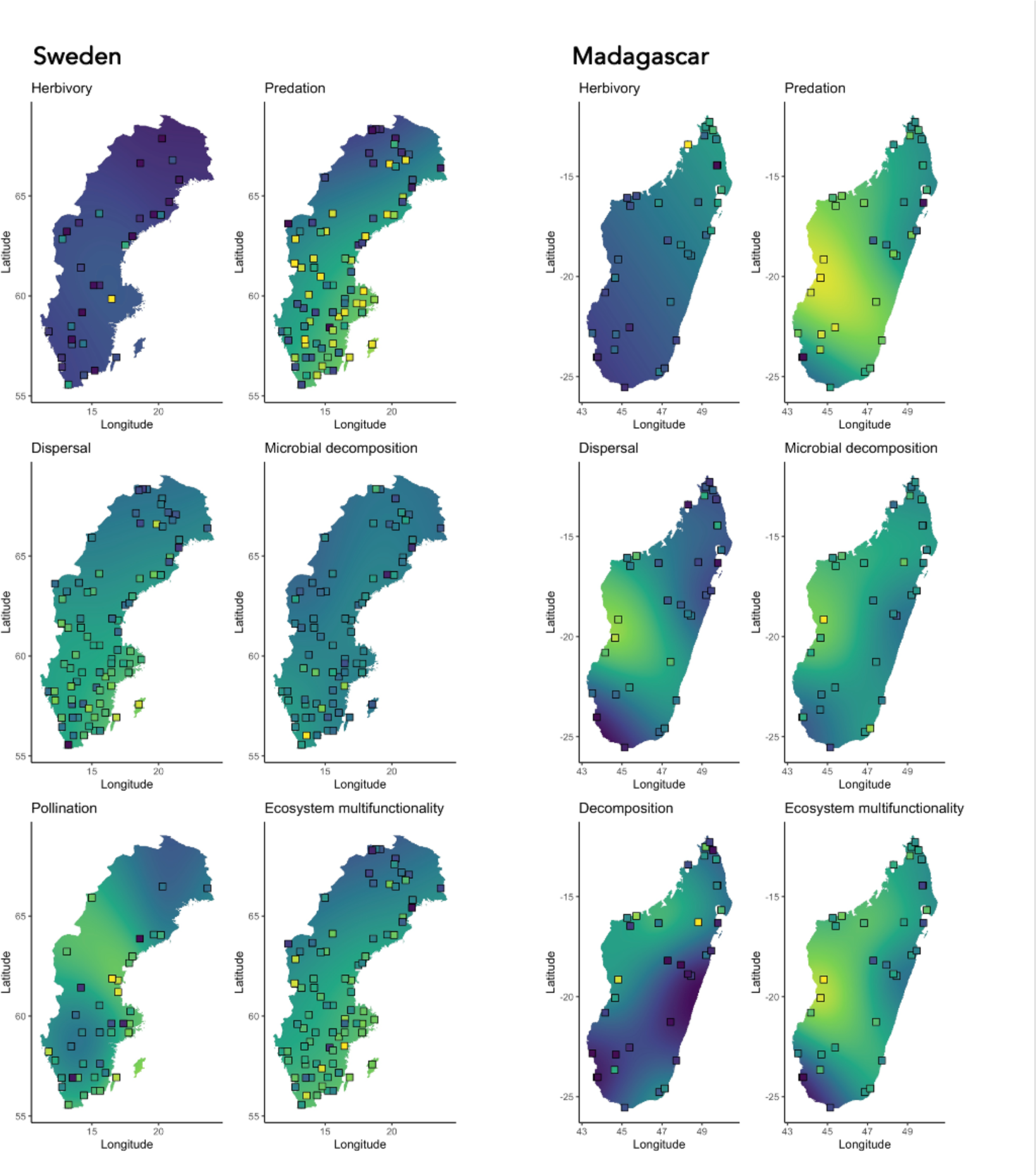
Rates of ecosystem functioning in Sweden (left) and Madagascar (right), for individual ecosystem functions (herbivory, predation, seed dispersal, pollination, microbial decomposition, and decomposition) as well as ecosystem multifunctionality, with locations of the experiments indicated with squares. Colors of the maps and colors of individual datapoints (squares) represent the relative functional rate, where lighter colors (yellow) indicate higher functional rates, and darker colors (blue) indicate lower functional rates. Functional rates are indicated as a color gradient on the background maps as based on smoothed predictions from generalized additive models with longitude and latitude as predictors.

### Ecosystem function experiments

To investigate the rate of ecosystem functioning, we conducted measurements across Sweden and Madagascar on a set of ecosystem functions: herbivory, seed dispersal, predation, decomposition and pollination (Sweden only) (Table S1). While conceptually targeting the same set of ecosystem functions, we note that practical considerations associated with working in the tropics (Madagascar) versus the temperate zone (Sweden), such as challenging logistics in Madagascar versus the availability of more solid infrastructure in Sweden, caused some differences in the methods employed. These differences concerned the specifics of how individual functions were measured, and the inclusion of an additional function (pollination) in Sweden, whereas the overall approach to measuring multifunctionality was the same across regions. For a detailed description of all ecosystem function experiments, see Text S1. For a visual overview of ecosystem function measurements, see Figure S3. For a summary of functions and methods measured per region, see Table S1.

In Sweden and Madagascar, **herbivory** was measured by recording the proportion of leaves eaten by insect herbivores. In Sweden, we recorded herbivory rates by placing 10 willow cuttings at each trap site (n = 72 sites, fig. 1). The percentage of herbivory on each leaf was recorded after 23 to 45 days. To measure herbivory rates in Madagascar, we laid out four transects of 10 meters in different directions from the trap (n = 50 traps, Fig. 1), and randomly picked 25 leaves along each transect. The percentage of herbivory on each leaf was visually estimated by the same observer.

In Sweden and Madagascar, **seed dispersal and predation** were measured by recording the proportion of removed seeds and prey during a set period of time. In Sweden, we placed three boxes with each 10 seeds of 10 different plant species and 1 gram of dried mealworms at each trap location (n = 171 traps, Fig. 1). The remaining number of seeds and mealworms were quantified after approx. 48 hours. In Madagascar, we filled 20 holder devices with 10 seeds from two different plant species, 20 holders with equal amounts of dried mealworms and crickets, and 20 holders with equal amounts of tinned sardines at each trap location (n = 50 traps, Fig. 1). Holders were placed along four transects of five meters in different directions from the Malaise trap. The remaining seeds and prey items were quantified after 90 minutes.

We measured **decomposition** rates in the soil in both Sweden and Madagascar. In Sweden we focused on microbial decomposition, while in Madagascar we additionally measured decomposition by soil-dwelling invertebrates. To measure decomposition by microorganisms, we used the teabag method in both Sweden (n = 163 traps, Fig. 1) and Madagascar (n = 50 traps, Fig. 1) (Keuskamp et al. 2013). Teabags were weighed after about 60 days in Madagascar and 90 days in Sweden. To measure decomposition by invertebrates in Madagascar, we additionally placed lamina baits of two different substrates (dried bean- and eucalyptus leaves) at each trap site (n = 50 traps, Fig. 1) (Kratz 1998). After three weeks, we assessed the proportion of substrate that remained.

We measured **pollination** rates in Sweden by planting four strawberry plants in soil bags at each site (n = 78 traps, Fig. 1). When matured, ripe strawberries were collected, and the total and fertilized number of seeds on each strawberry were counted to obtain the fertilization rate. Since all trap sites in Madagascar were placed in forests where flowering plants are less common, we only measured pollination at the trap sites in Sweden.

### Abiotic and biotic drivers

To quantify abiotic impacts on ecosystem functioning, we obtained data on climate and landscape characteristics (for methodological details, see Text S2 and Table S2). We collected data on temperature and vegetation cover from the ERA5-land database. Soil moisture and leaf litter were measured at each site, at five locations around the trap.

To investigate whether ecosystem functioning was driven by biotic factors, i.e., insect communities, we set up a Malaise trap at each experimental site (Fig. S3, Text S2). We obtained insect biomass, species richness and community composition as metrics for the biotic community. Insect biomass was wet-weighed for each sample, from which we calculated the mean insect biomass per day per trap. To uncover the insect species present in each catch as well as the biomass of insects in each sample, Malaise trap catches were processed and metabarcoded as described in Iwaszkiewicz-Eggebrecht (2023) (Text S2). We used clusters as a proxy for species for the purpose of this manuscript. Species richness was calculated as the mean species richness per day for each trap. As a univariate metric of community composition, we obtained the first main coordinate axis from a principal coordinates analysis (PCoA) on a Jaccard distance matrix (function *vegdist* in the *vegan* package, R v.4.2.0), with species presence or absence records at the trap level.

### Statistical analyses

#### Drivers behind ecosystem functioning

To analyse the direct and indirect drivers of ecosystem functioning in tropical and temperate zones, we fitted structural equation models (SEM) including abiotic and biotic factors (Table S3). Structural equation modelling allowed us to analyse the causal structure of the drivers behind ecosystem functioning, enabling us to not only look at direct effects of abiotic and biotic drivers, but also at mediating variables contributing to ecosystem functioning. As climatic and landscape drivers, we included temperature, soil moisture, vegetation cover and leaf litter depth, and as biotic drivers, we included several descriptors of the local insect community, including species richness, total insect biomass and community composition.

We constructed initial conceptual models based on a priori knowledge about potential direct and indirect drivers behind ecosystem functioning, with a separate model for Sweden (Fig. S4a-c) and Madagascar (Fig. S5a-c). Conceptual models included the hypothesized relationships between climate, landscape, biotic community and ecosystem functions. In these models, we included all climate and landscape variables as predictors for the biotic community, and the biotic community as a predictor for ecosystem functions (the biotic metrics of species richness, biomass and community composition were analysed in separate models, see Table S3 and Fig. S4 for model structures). To capture direct relations between climate, landscape and ecosystem functions, temperature and soil moisture were included as direct drivers behind all ecosystem functions, whereas vegetation cover was included as a direct driver for aboveground ecosystem functions (i.e. herbivory, seed dispersal, predation and pollination) and leaf litter for belowground ecosystem functions (i.e. decomposition by invertebrates and microbes). Structural equation models were fitted using the *psem* function in the package *PiecewiseSEM* (Lefcheck 2016). All predictor variables were scaled to zero mean and unit variance. To obtain model estimates and statistical significance of direct, indirect and total effects on ecosystem functioning of all predictors in the model, we used the summary function with manual calculation of path coefficients for indirect effects (Lefcheck 2016).

#### Drivers behind ecosystem multifunctionality

To investigate the drivers behind ecosystem multifunctionality, we constructed a structural equation model including all abiotic and biotic drivers (Table S4, Fig. S4d-f, Fig. S5d-f). To obtain a metric of ecosystem multifunctionality for Sweden and Madagascar, we used the averaging approach (Maestre et al. 2012): We calculated the z-value for each ecosystem function at a site, and then obtained the mean z-score across functions for each site, separately for Sweden and Madagascar. To prevent bias towards certain functions in our multifunctionality metric, we averaged the z-scores of the two metrics for decomposition before averaging across functions.

#### Raw and residual correlations among ecosystem functions

To investigate whether individual ecosystem functions were positively or negatively correlated with each other at the site level, we fitted raw correlations across ecosystem functions using a Pearson correlation test (function *cor.test*), separately for Sweden and Madagascar. Here, a negative correlation reflects a trade-off between functions, whereas positive correlations indicate that ecosystem functions are varying in concert.

To establish whether correlations between individual ecosystem functions were caused by individual ecosystem functions responding similarly to the same drivers, or whether additional associations remained after accounting for these environmental imprints, we i) used the SEM analyses to infer the main structuring forces across ecosystem functions, both for Sweden and Madagascar; ii) fitted simple linear models to these ecosystem functions, with the factors identified as covariates (Table S5); iii) extracted the residuals and iv) checked for remaining residual correlations. To identify the individual contribution of abiotic (climate and landscape) and biotic variables in explaining correlation structures among functions, we also fitted separate linear models for abiotic variables (climate and landscape) and biotic variables, of which we extracted residuals and checked for residual correlations. Residual correlations among ecosystem functions may suggest either remaining associations among functions, or the existence of joint hidden drivers not accounted for.

## Results

### Drivers behind ecosystem functioning

In Sweden, the main driver behind ecosystem functioning was temperature, which had a strong direct positive effect on seed dispersal, predation, and microbial decomposition (Fig. 2a, Table S6). Moreover, sites with more vegetation cover had lower pollination rates, and tended to have less predation, compared to sites with less vegetation cover (Table S6). Temperature also affected the biotic community, with positive effects on insect richness and biomass, and a strong influence on the composition of communities (Table S6). Vegetation cover positively affected species richness, tended to positively affect insect biomass, and influenced community composition (Table S6). Community composition was the only biotic metric that significantly affected ecosystem functioning, affecting only predation rates (Table S6). Besides direct effects, temperature and vegetation cover also indirectly significantly affected predation rates, via changes in community composition (standardized coefficients = - 0.38 and -0.07, respectively, Fig. 1a, Table S6).

**Figure 2.**
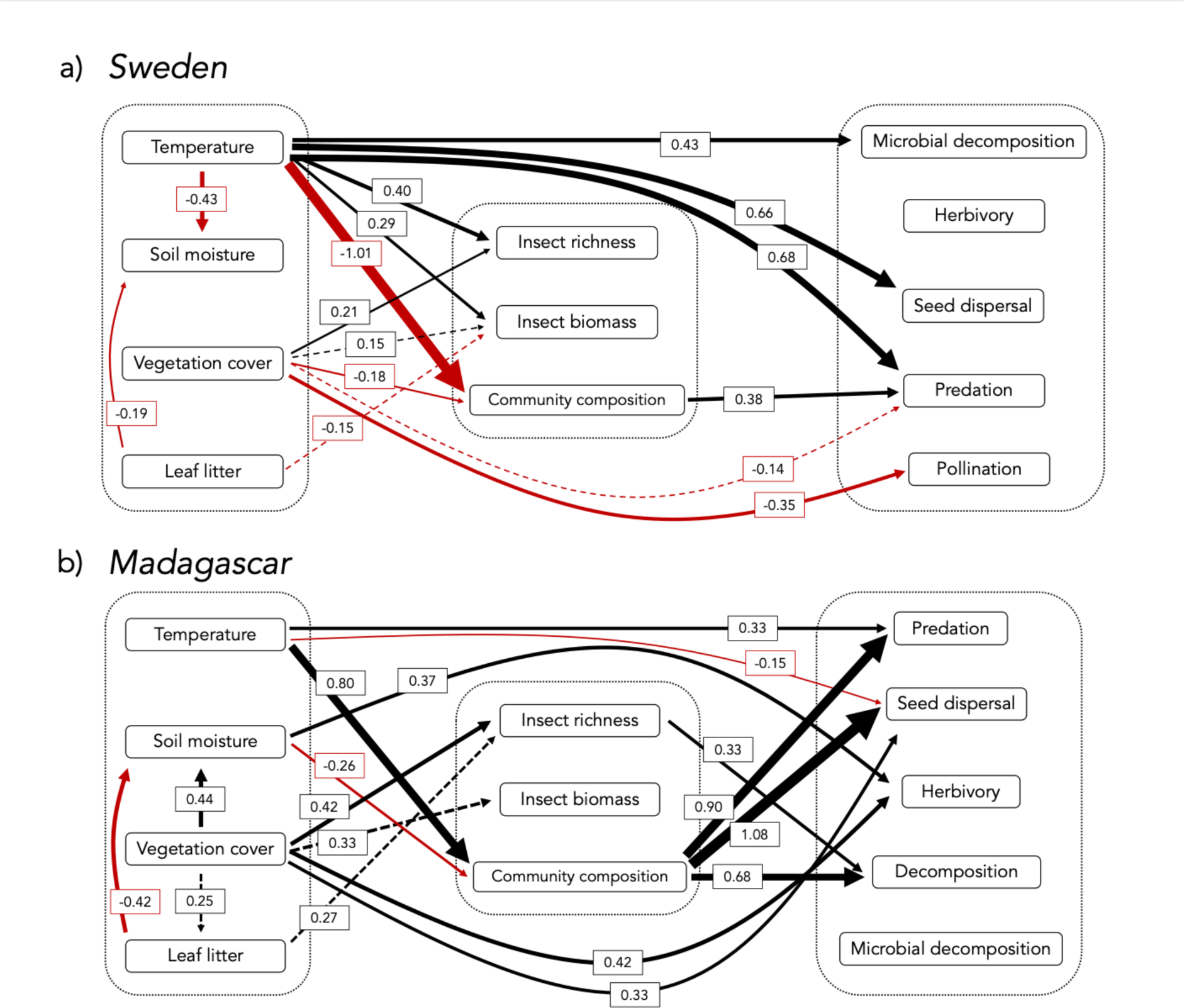
The effects of climate, landscape and biotic community on individual ecosystem functions in a) Sweden and b) Madagascar. Black arrows indicate positive effects, and red arrows indicate negative effects. Solid arrows indicate significant effects (p < 0.05), and dashed arrows indicate trends (0.05 < p < 0.10). For conceptual models, see Figs. S4a-c and S5a-c.

In Madagascar, temperature positively affected predation rates but negatively affected seed dispersal (Fig. 2b, Table S7). Vegetation cover and soil moisture had a direct, positive effect on herbivory (Table S7). Temperature and soil moisture influenced the composition of insect communities across sites (Table S7). Moreover, sites with more vegetation cover had higher insect richness, and tended to have more insect biomass (Table S7). Sites with more leaf litter tended to have higher species richness (Table S7). Of the biotic metrics, community composition had a strong impact on half of the measured ecosystem functions, including predation, seed dispersal and decomposition. Also, sites with higher insect species richness showed faster decomposition rates (Table S7). Besides direct effects, temperature and soil moisture also indirectly significantly affected predation (standardized coefficients = 0.72 and -0.23, resp.), seed dispersal (standardized coefficients = 0.86 and -0.28, resp.) and decomposition (standardized coefficients = 0.54 and -0.18, resp.) via effects on community composition (Table S7). Vegetation cover also indirectly significantly increased decomposition rates, by promoting insect richness (standardized coefficient = 0.14, Table S7).

### Drivers behind ecosystem multifunctionality

In Sweden, temperature had a strong positive effect on ecosystem multifunctionality, while vegetation cover had a negative effect (Fig. 3a, Table S8). Of the biotic metrics, community composition determined ecosystem multifunctionality rates, while species richness and biomass had no effect (Table S8). Via impacts on community compositions, temperature and vegetation cover also indirectly significantly affected ecosystem multifunctionality (standardized coefficients = -0.25 and -0.05 respectively, Fig. 3a, Table S8).

**Figure 3.**
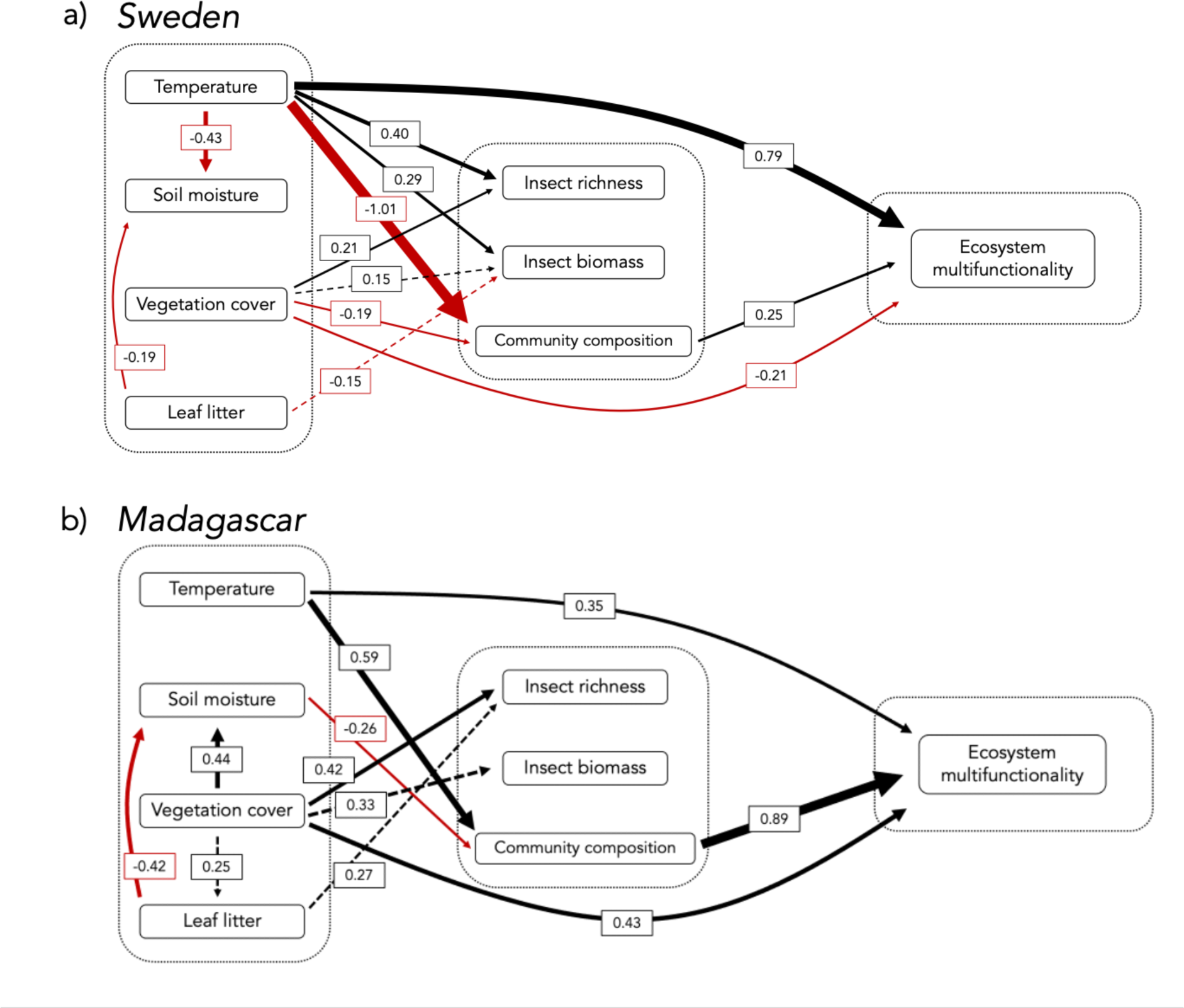
The effects of climate, landscape and biotic community on ecosystem multifunctionality in a) Sweden and b) Madagascar. Black arrows indicate positive effects, and red arrows indicate negative effects. Solid arrows indicate significant effects (p < 0.05), and dashed arrows indicate trends (0.05 < p < 0.10). For conceptual models, see Figs. S4d-f and S5d-f.

In Madagascar, temperature and vegetation cover positively affected ecosystem multifunctionality (Fig. 3b, Table S9), and differences in community composition across sites had a strong influence on ecosystem multifunctionality (Table S9). By influencing community composition, temperature and soil moisture indirectly significantly affected ecosystem multifunctionality (standardized coefficients = 0.53 and -0.23 respectively, Fig. 3b, Table S9).

### Correlations among ecosystem functions

In Sweden, sites with high seed dispersal rates also had high predation rates, while none of the other ecosystem functions showed positive or negative correlations among sites (Fig. 4a). The positive correlation between seed dispersal and predation remained after taking into account the effect of climate, landscape and biotic community (Fig. 5a).

**Figure 4.**
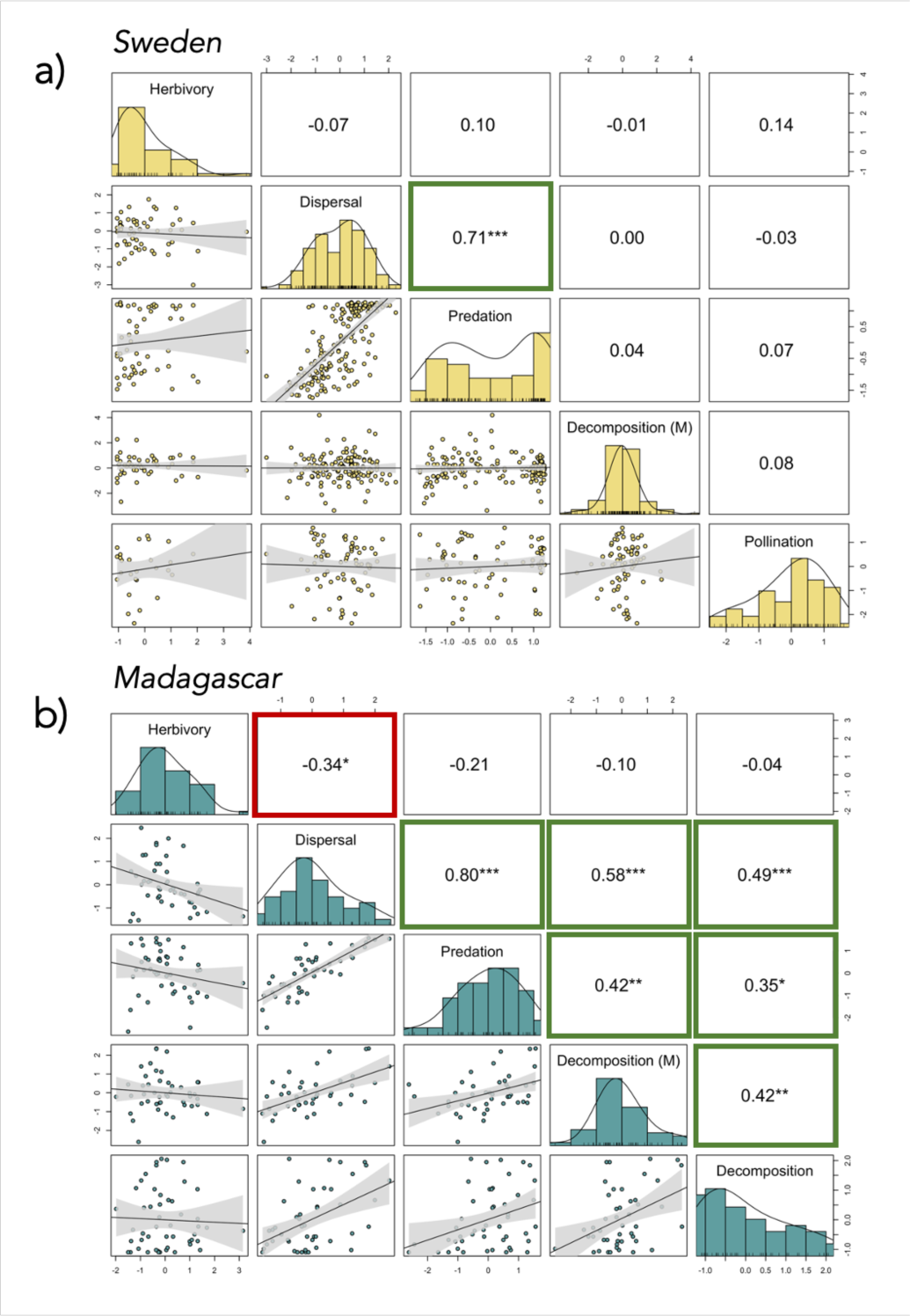
Raw correlations among ecosystem functions in Sweden (a) and Madagascar (b). For Sweden, ecosystem functions include herbivory, seed dispersal, predation, microbial decomposition (“M”) and pollination. For Madagascar, ecosystem functions include herbivory, seed dispersal, predation, microbial decomposition (“M”) and decomposition by invertebrates and microbes. Linear correlations between ecosystem functions are presented in the lower left corner of the panels, and the Pearson correlation coefficient (r) with stars of significance (* p < 0.05, ** p < 0.01, *** p < 0.001) in the upper right corner of the panels. Green-bordered squares indicate significantly positive correlations, and red-bordered squares indicate significantly negative correlations. The histograms in the diagonal present the distribution of the data for all ecosystem functions.

**Figure 5.**
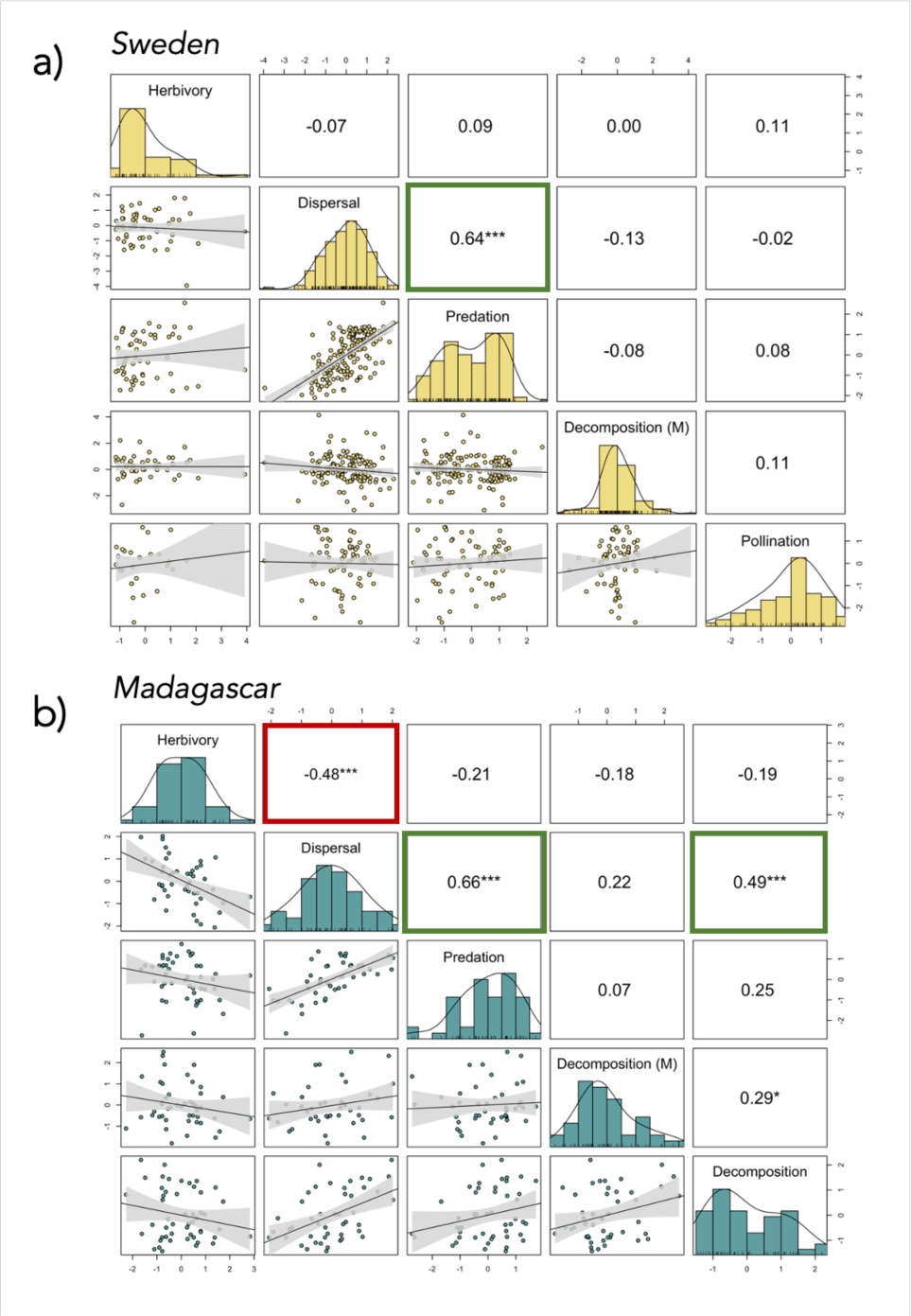
Residual correlations among ecosystem functions in Sweden (a) and Madagascar (b), after accounting for the main structuring forces identified by SEMs. For Sweden, ecosystem functions include herbivory, seed dispersal, predation, microbial decomposition (“M”) and pollination. For Madagascar, ecosystem functions include herbivory, seed dispersal, predation, microbial decomposition (“M”) and decomposition by invertebrates and microbes. Linear correlations between ecosystem functions are presented in the lower left corner of the panels, and the Pearson correlation coefficient (r) with stars of significance (* p < 0.05, ** p < 0.01, *** p < 0.001) in the upper right corner of the panels. Green-bordered squares indicate significantly positive correlations, and red-bordered squares indicate significantly negative correlations. The histograms in the diagonal present the distribution of the data for all ecosystem functions. For residual correlations structures of ecosystem functions in Madagascar after accounting for abiotic and biotic variables separately, see Fig. S6.

In Madagascar, all measured ecosystem functions except herbivory — i.e., seed dispersal rates, predation rates, microbial decomposition and decomposition by microbes and invertebrates — were positively correlated across sites (Fig. 4b). Sites with high herbivory had lower seed dispersal rates, but no other ecosystem functions correlated to herbivory. However, with climatic, landscape and biotic effects accounted for, most of the positive correlations among functions disappeared (4 out of 6 positive correlations, Fig. 5b), with only the positive correlation between seed dispersal and predation and seed dispersal and decomposition remaining. The negative correlation between herbivory and seed dispersal also remained after accounting for climatic, landscape and biotic effects (Fig. 5b). Abiotic (climate and landscape) and biotic conditions contributed equally to drive positive correlations among functions (Fig. S6). However, negative correlations among herbivory and predation, as well as between herbivory and microbial decomposition, appeared only after taking into account abiotic conditions (Fig. S6a).

## Discussion

In this study, we aimed to pinpoint the abiotic and biotic drivers behind ecosystem functioning and multifunctioning in a temperate and a tropical climate, and to discriminate potential differences in strength between these biomes. We were particularly interested in whether ecosystem functions increase in concert, or trade off against each other. In Sweden, we found a strong effect of temperature on ecosystem functionality and multifunctionality, with other factors, especially biotic ones, exerting much weaker effects. In the tropical biome of Madagascar, effects of biotic drivers were much stronger than effects of abiotic drivers. These findings support general theory, positing that biotic interactions will be stronger in tropical than temperate zones (Roslin et al. 2017, Paquette and Hargreaves 2021), thus accentuating the relative role of climate at higher latitudes. Following this reasoning, theory predicts that climate would drive positive correlations among functions in Sweden, whereas biota would drive positive correlations among functions in Madagascar. Nonetheless, our results revealed a general absence of correlations among functions in Sweden, and the structure of these correlations was largely independent of abiotic or biotic drivers. In Madagascar, all functions except herbivory positively correlated with each other in space, and most of these correlations could be explained by accounting for local climate, landscape features and biota.

### Temperature is the main driver of ecosystem functioning in the temperate but not the tropical region

Temperature emerged as a key driver of functionality in Sweden, while in Madagascar, climate or landscape showed no more pronounced effect on ecosystem functioning than the other. As expected, warmer sites in Sweden sustained higher functionality rates of individual functions as well as higher levels of multifunctionality. In Madagascar, predation rates as well as multifunctionality rates were positively affected by temperature, whereas temperature did not stand out as the dominant driver of ecosystem functioning, and even had a slight negative effect on seed dispersal rates.

The difference between tropical Madagascar and temperate Sweden may perhaps be attributed to differences in the mean level of energy available in the two systems. At higher latitudes, the activity of organisms and chemical processes is likely temperature-limited for much of the year, whereas in tropical regions the temperatures are more often high enough to sustain such activities. Supporting this finding, Swedish sites with more vegetation had reduced pollination, predation, and multifunctionality rates, while Malagasy sites with more vegetation exhibited increased rates of seed dispersal and herbivory, as well as higher multifunctionality rates. This is consistent with differences in shading impacts against different average temperatures. In Sweden, blocking of solar radiation by vegetation cover prevents local environments from warming and may result in reduced functionality. In a tropical climate like Madagascar, protection by vegetation cover against solar radiation may be beneficial for ecosystem functionality by reducing the risk of extremely hot temperatures and droughts (Fetcher et al. 1985).

As a further illustration of different abiotic impacts in response to different average climatic conditions, we found that soil moisture had no effect on functionality rates in Sweden, whereas in Madagascar, moister sites experienced higher herbivory rates. This contrast likely reflects the more limited availability of water in the tropics, especially in dry forests. Plants growing in moister areas are likely to be more attractive to herbivores than plants growing in dry forests, where sclerophyllous plants and spiny bushes are the dominant vegetation types.

Considering that both the climate and the landscape are undergoing major changes due to anthropogenic activities, revealing the impact of climate and landscape on the functioning of our ecosystems across zones is of vital importance. To do so, we need further studies conducted across multiple countries in both zones to confirm potential differences in the impact of climate and landscape across tropical and temperate biomes.

### Biodiversity had more pronounced effects on ecosystem functioning in the tropical region than the temperate region

Overall, we detected few significant effects of biodiversity on functioning in the temperate zone, but found more pronounced effects of biodiversity in the tropics. In the temperate zone, community composition as scored by Malaise traps had an effect on predation as well as on ecosystem multifunctionality rates, even though effect sizes were relatively small. In the tropics, community composition strongly affected predation, seed dispersal and decomposition, and was the strongest driver of multifunctionality rates. Moreover, richness had a positive effect on decomposition in the tropics, but no effect on functioning in the temperate zone. Hence, our findings suggest that effects of biodiversity on functioning are dependent on the biome of the ecosystem, thus hampering generalization regarding biodiversity-functioning relationships in natural contexts (van der Plas 2019, Hagan et al. 2021). While the biodiversity-functioning relationship in experimental settings is most often found to be positive (Tilman et al. 2014), species assemblages are pre-selected based on certain traits, and hence, the effects of biodiversity on functioning may often be overestimated. Our findings also highlight that effects of biodiversity are strongly dependent on the metric of biodiversity adopted: While species biomass and richness had zero to minimal effects on functioning, the composition of communities proved to be an important driver, suggesting that what matters is not the *amount* or *number* of species per se, but rather the *identity* of species present in the community (Mahaut et al. 2020). Thus, to uncover general patterns of the effects of biodiversity on functioning in real-world communities, we urgently need further studies conducted at large spatial scales, exploring different aspects of biodiversity, which will enable us to untangle the effects of different biotic drivers on functioning of ecosystems across climatic zones.

Besides the direct effects of biodiversity, biodiversity also mediated some of the effects of climate and landscape on ecosystem functioning in both Sweden and Madagascar. In Sweden, temperature and vegetation cover affected community composition, which in turn affected predation and ecosystem multifunctionality. In Madagascar, temperature and soil moisture affected community composition, which in turn affected predation, seed dispersal, decomposition and ecosystem multifunctionality. Vegetation cover also indirectly increased decomposition rates by promoting species richness. While mediating effects of biodiversity are often neglected in current studies, we argue that this is an important component to consider when aiming to understand the full extent of effects that biodiversity has on the functioning of natural communities (Hooper et al. 2005, Hisano et al. 2018). Moreover, future studies could investigate whether biodiversity can buffer potentially negative effects of climate change and land-use changes on ecosystem functioning across biomes.

### More functions increased in concert in Madagascar than in Sweden

Based on harsher climatic conditions and poorer insect communities imposing more limits on ecosystem functioning, we expected that functions in the temperate region would be more tightly interlinked than functions in the tropical region (Fig. S1). Nonetheless, we found the opposite pattern: Functions were more positively correlated with each other in Madagascar than in Sweden. In Madagascar, all functions were positively correlated except herbivory, and most of these positive correlations in Madagascar could be explained by climate, landscape and local biota. These results suggest that sites in the tropics tend to be either “highly functional” or “poorly functional” across functions, and that local climate, landscape features and local biota largely determine functionality rates across functions at a site. Still, some correlations (3 out of 7) remained after accounting for the abiotic and biotic environment. Thus, the remaining positive and negative correlations among functions in the tropics are likely driven by an environmental or biotic variable unaccounted for by our study. Pinpointing these additional drivers thus emerges as a key priority for future studies, as the current study design did not allow us to resolve all drivers of correlated functionality.

## Conclusion

Our study achieved to describe the patterns and drivers of ecosystem functioning at an unprecedented scale, with intense spatial replication across two major countries, one in the tropical and one in the temperate zone. While awaiting global sampling to confirm the generality of our findings, this study sets the baseline expectation that patterns and drivers of ecosystem functioning fundamentally differ between the temperate and tropical zone: Functions were highly correlated in space in the tropics but not the temperate zone, and, while temperature was the main driver of functioning in the temperate zone, the biotic community was the most important driver in the tropics. Resolving such differences among biomes will be key to correctly predict shifts in ecosystem functioning in response to environmental disturbances.

## Supplementary

**Text S1.** Methodological details on ecosystem function experiments conducted in Sweden and Madagascar.

*In Sweden and Madagascar, **herbivory** was measured by recording the proportions of leaves eaten by insect herbivores. In Sweden, we recorded herbivory rates by placing willow cuttings close to a Malaise trap (n = 72 traps, fig. 1). Willow was chosen as the focal plant since it is clonal (thus excluding effects of genetic heterogeneity), easy to propagate, and a widely used host plant for insects. For this, we placed a bag with 15 litres of potting soil 5 meters from the back of the Malaise trap, slicing 10 openings across the bag, in which we planted 10 willow cuttings. The soil bag was cut from each of four sides to allow water drainage while retaining some moisture, and the bag was thoroughly watered after the willow sticks were planted. Willow cuttings remained in the field for 23 to 45 days, after which the percentage of herbivory on each leaf was recorded. To measure herbivory rates in Madagascar, we laid out four transects of 10 meters in different directions from the trap (n = 50 traps, Fig. 1), and randomly picked one leaf from high vegetation and one leaf from low vegetation approximately every meter, for a total of 25 leaves per transect, and 100 leaves per trap location. Directly after collection, leaves were dried in a plant press. After drying, the percentage of herbivory on each leaf was visually estimated by the same observer*.

*In Sweden and Madagascar, **seed dispersal and predation** were measured by recording the proportion of removed seeds and prey during a set period of time. Specifically, in Sweden, at each trap location (n = 171 traps, Fig. 1), we filled three boxes with sand, and added 10 seeds of 10 different plant species to each box (Helianthus annuus, Avena sativa, Lolium perenne, Linum usitatissimum, Anemone nemorosa, Sesamum indicum, Plantago lanceolata, Trifolium pretense, Viola tricolor and Papaver somniferum), resulting in 100 seeds per box. To measure predation rates, we also added 1 gram of dried mealworms to each box, equivalent to 20 mealworms based on average mealworm weight (0.05 grams). We ensured that seeds and mealworms were spread evenly across the sand surface. The boxes were then closed with a lid. Six holes of 1 cm diameter on the side of each box allowed arthropods access to the box. This setup excluded other animals that feed on seeds or worms, such as birds and rodents. Boxes were secured to the ground with pegs and placed 2.5 meters away from a Malaise trap. Boxes were collected from the field after 2 days (approximately 48 hours), and the remaining number of seeds and mealworms were counted. To measure seed dispersal and seed predation in Madagascar at each trap location (n = 50 traps, Fig. 1), we filled 20 holder devices (circular plastic dish with an opening in the middle to allow attachment to the ground) with 10 seeds from two different plant species (sunflower and sesame). Additionally, to measure predation rates, we filled 20 holders with 3 dried mealworms cut in half and five dried crickets, and 20 more holders with 1.4 to 1.8 grams of tinned sardines, from which excessive oil was removed with toilet paper before weighing and placement. We laid out four transects of five meters in different directions from the Malaise trap, and placed one holder with seeds, one with mealworms and crickets, and one with sardines at every meter along each of the transects, resulting in 5 holders of each type along each transect, for 15 holders per transect in total. Holders were secured to the ground with a peg through the middle of the device. Each holder was individually covered with a lid to exclude animals other than arthropods. After 90 minutes, we counted the remaining seeds and prey items and weighed the remaining sardines*.

*We measured **decomposition** rates in the soil in both Sweden and Madagascar. In Sweden we focused on microbial decomposition, while in Madagascar we additionally measured decomposition by soil-dwelling invertebrates. To measure decomposition by microorganisms, we used the teabag method in both Sweden (n = 163 traps, Fig. 1) and Madagascar (n = 50 traps, Fig. 1) (Keuskamp et al. 2013). With a soil corer, we made two adjacent holes 8 cm deep in the soil, 15 cm apart, at three different locations surrounding the trap, resulting in six holes at each trap. At each pair of holes, we placed one teabag filled with green tea, and one teabag filled with red tea (rooibos), and covered the bags with soil. Teabags were left in the soil to decompose for about 60 days in Madagascar and 90 days in Sweden, then collected, stored in silica gel, and weighed. To measure decomposition by invertebrates in Madagascar, we additionally placed lamina baits of two different substrates (dried bean- and eucalyptus leaves) around the Malaise traps (n = 50 traps, Fig. 1) (Kratz 1998). With a soil corer, we made three 10 cm-deep holes in the soil (one per teabag location, 15 cm apart), and in each hole, we placed one lamina bait of each substrate-type 3– 5 cm from each other. Lamina baits were left in the soil for three weeks to decompose, after which baits were visually inspected by one observer to assess the proportion of substrate (scored as 0, 0.5 or 1) that remained*.

*We measured **pollination** rates in Sweden by recording the proportion of fertilized seeds of strawberry plants placed in the field. Since all trap sites in Madagascar were placed in forests where flowering plants are less common, we only measured pollination at the trap sites in Sweden (n = 78 traps, Fig. 1). Specifically, we used a clonal version of the German woodland strawberry (Fragaria vesca) cultivar “Rügen” produced by Mälarö Odling AB (Ekerö, Sweden). We chose this cultivar because it is everbearing, produces flowers throughout the summer, and is mostly dependent on insects for pollination (Evans and Jones 1967). We cut four holes in the corners of the same soil bags as used for the willow sticks (placed at 5 meters behind each Malaise trap), and planted strawberry plants there. Any flowers already present at the time of planting were removed, ensuring that flowers only experienced pollination at the trap site. Plants were left in the field until the strawberries matured. Ripe strawberries were collected, and the total and fertilized number of seeds on each strawberry were counted to obtain the fertilization rate*.

**Text S2.** Methodological details on the collection of abiotic and biotic variables in Sweden and Madagascar.

*To quantify abiotic impacts on ecosystem functioning, including effects of climate and landscape characteristics, we obtained information on temperature, soil moisture, leaf litter and vegetation cover at each experimental site in Sweden and Madagascar (Table S2). To obtain information on temperature, temperature data at 2 meters height was downloaded from the ERA5-Land database at a spatial resolution of 11 km^2^, and then downscaled to 2 km^2^ using elevation as a covariate (Kusch and Davy 2022). Soil moisture was measured with the SM150 Soil moisture kit (DeltaT Devices Ltd., UK) at five locations around the experimental site (Fig. S3). Leaf litter depth was measured with a ruler at five locations around the experimental site (Fig. S3). Information on vegetation cover was obtained from ERA5-Land, where we downloaded data on the leaf area index (LAI) of high vegetation (m^2^ m^-2^) within an area of 11 km^2^ around the experimental site. In Sweden, ecosystem function experiments were conducted over the course of the growing season. Hence, for the climate (temperature) and landscape (vegetation cover) predictors, we used the site-level mean values of each predictor over the entire growing season (May – September) in our models (Table S2). In Madagascar, ecosystem functions were measured throughout the year, excluding the dry season when no experiments were conducted. In this case, we used the site-level mean values of temperature and vegetation cover during a two-month period surrounding the experimental date (Table S2)*.

*To investigate whether ecosystem functioning was driven by biotic factors, i.e., insect communities, we set up a Malaise trap at each experimental site (Fig. S3). To uncover the insect species present in each catch as well as the biomass of insects in each sample, Malaise trap catches were processed and metabarcoded as described in Iwaszkiewicz-Eggebrecht (2023). The arthropod biomass from each catch was assessed by wet-weighing, then the samples were subjected to a mild lysis and DNA was extracted from the lysate. Subsequently, libraries targeting a 418-bp fragment of a COI barcode were prepared through a two-step PCR procedure. All metabarcoding libraries were sequenced on an Illumina NovaSeq 6000 SPrime platform at a depth of approximately 1M reads per sample and processed bioinformatically following pipelines for read trimming and filtering (https://github.com/biodiversitydata-se/amplicon-multi-cutadapt) and barcode reconstruction and taxonomic annotation (https://nf-co.re/ampliseq) using a custom-made reference COI database (https://doi.org/10.17044/scilifelab.20514192.v4). We removed ASVs that were chimera’s, ASVs that had STOP codons shifting the reading frame of the gene, ASV that had deletions shifting the reading frame of the gene, ASVs present in more than 5% of blanks, and ASVs with less than three counts summed across the dataset. After filtering, we recovered 540.665 unique ASVs for Sweden and 544.405 unique ASVs for Madagascar. We then clustered those based on sequence similarity using swarm, with the threshold for clustering at 13. We identified 70.293 and 168.299 clusters in Sweden and Madagascar respectively. We used clusters as a proxy for species for the purpose of this manuscript. From this data, we obtained insect biomass, species richness and community composition as metrics for the biotic community. In Sweden, we only used Malaise trap samples collected from May to September (approx. 10 samples per trap, Table S2). In Madagascar, we used trap samples collected during the 3 months surrounding the experimental date (1.5 month before and 1.5 month after the experiment, approx. 13 samples per trap, Table S2). Species richness was calculated at the sample level, and corrected for the number of sampling days (sum of unique species per sample/number of collection days), from which we calculated the mean species richness per day for each trap. Insect biomass was weighed for each sample, and divided by the number of collection days, from which the mean insect biomass per day per trap was calculated. To obtain a univariate metric of community composition at each site, we conducted a principal coordinates analysis (PCoA) on a Jaccard distance matrix (function vegdist in the vegan package, R v.4.2.0) including all recorded species, with their presence or absence records at the trap level. From the fitted PCoA, we used the first main coordinate axis as a proxy for differences in community composition across trap sites*.

*To characterise general biotic diversity, we opted to sample the most species-rich part of the local fauna, i.e. flying insect communities, using Malaise traps. These samples capture local variation in the most speciose part of communities, and thus reflect overall features of the biotic environment. Of these samples, we characterised several different dimensions of diversity, i.e. species richness, biomass and community composition. At the same time, it will be evident that the overall insect community will consist of widely different functional groups, ranging from pollinators to saprophages. Thus, different components of the summary community will – by default – contribute unequally to individual functions, and the overall metrics of the insect community may be poorly reflective of the species richness, biomass or composition of the actual fauna contributing to our metrics of ecosystem functioning. The indirect nature of the biotic measures we employed should be kept in mind when gauging the relative impact of the biotic drivers. At the same time, given the difficulties in dissecting the fauna into functional guilds, in particular for Madagascar, and the very fact that holometabolous insects will typically switch between guilds during their life cycle, we believe that our overall metrics provide a fair proxy for biotic impacts. This applies in particular to ecosystem multifunctionality. For this measure, we should logically focus on the overall features of the biotic community rather than any specific guild or taxon*.

**Table S1.**
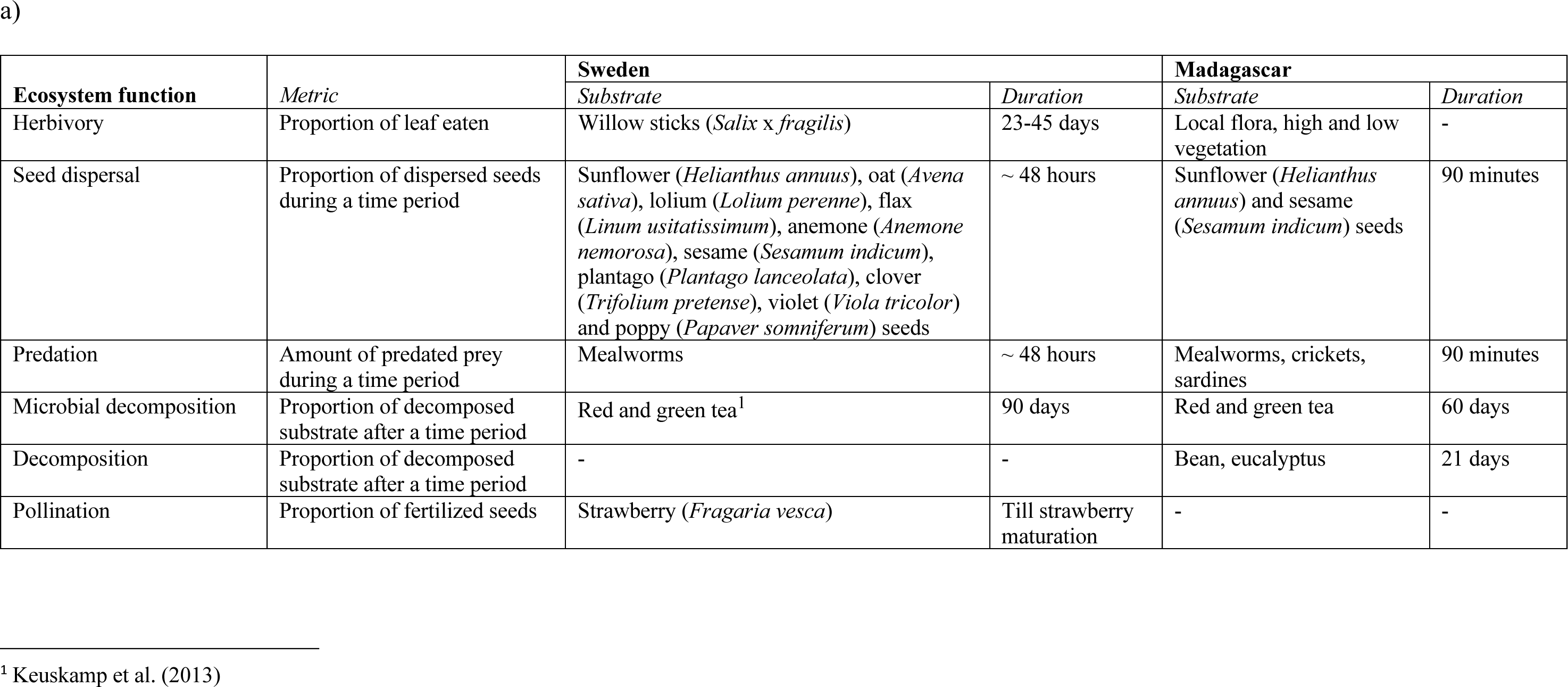
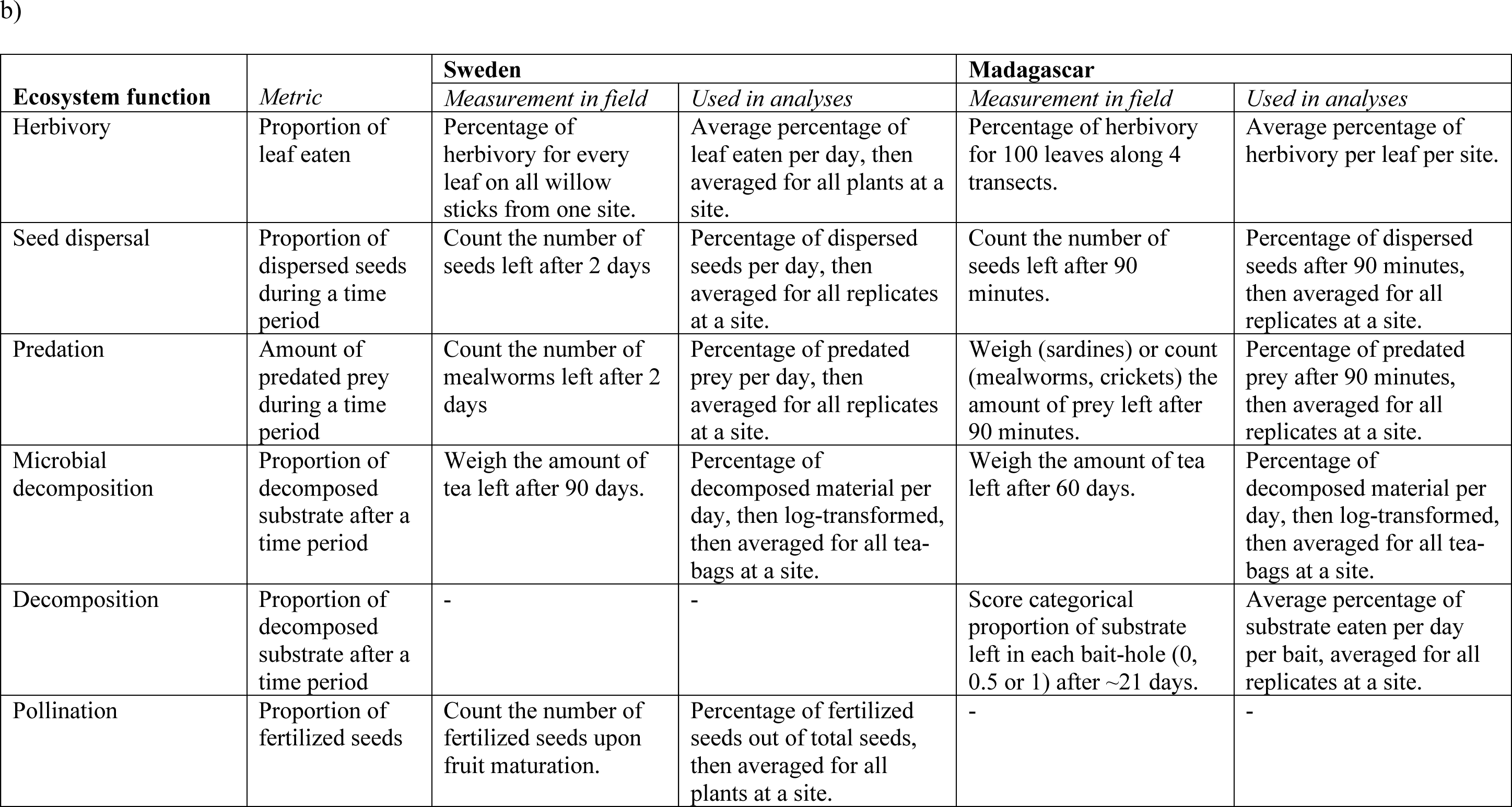
Experiments and metrics of ecosystem functioning in Sweden and Madagascar. Panel a) shows an overview of the experiments conducted on ecosystem functioning in Sweden and Madagascar. In Sweden, we conducted experiments on seed dispersal, predation, herbivory, microbial decomposition and pollination. In Madagascar, we conducted experiments on seed dispersal, predation, herbivory, microbial and invertebrate decomposition. Duration of the experiments were adapted to local conditions. Panel b) shows an overview of the translation of field measurements into metrics for data analysis. For a visual schematic overview of all ecosystem function measurements, see Figure S2.

**Table S2.** Overview of climatic, landscape and biotic drivers used in the analysis. For more details on the experimental design of on-site measurements, see Figure S2.

| Driver | Method | Time period Sweden | Time period Madagascar |
| --- | --- | --- | --- |
| *Abiotic drivers* |  |  |  |
| Temperature | Data on temperature (2 m height) downloaded from the ERA5-Land database, with a spatial resolution of 11 km<sup>2</sup>, then downscaled to 2 km<sup>2</sup>. | Downloaded data during the growing season, from May to September. | Downloaded data within a 2-month interval surrounding the experimental date, i.e., 1 month before the experiment, and 1 month after the experiment. |
| Soil moisture | On-site measurements with SM150 soil moisture kit, at 5 locations around the experimental site. | Measured when long-term ecosystem function experiments were set up (pollination, herbivory, and decomposition). | Measured when ecosystem function experiments were conducted. |
| Leaf litter | On-site measurements with ruler, at 5 locations around experimental site. | Measured when long-term ecosystem function experiments were set up (pollination, herbivory, and decomposition). | Measured when ecosystem function experiments were conducted. |
| Vegetation cover | Data on leaf area index (LAI) of high vegetation downloaded from the ERA5-Land database, with a spatial resolution of 11 km<sup>2</sup>. | Downloaded data during the growing season, from May to September. | Downloaded data within a 2-month interval surrounding the experimental date, i.e., 1 month before the experiment, and 1 month after the experiment. |
| *Biotic drivers* |  |  |  |
| Species richness | Species richness in each sample, corrected for sampling days, from which mean species richness per day per trap was calculated. | Samples collected during the growing season (May – September) | Samples collected during 3 months surrounding the experimental date (1.5 month before and 1.5 month after). |
| Community composition | PCA on matrix of all present and absent species in each trap. | Samples collected during the growing season (May – September) | Samples collected during 3 months surrounding the experimental date (1.5 month before and 1.5 month after). |
| Insect biomass | Wet-weighing of insect biomass in Malaise-trap samples. | Samples collected during the growing season (May – September) | Samples collected during 3 months surrounding the experimental date (1.5 month before and 1.5 month after). |

**Table S3.** Overview of fitted structural equation models to analyze the direct and indirect effects, i.e., via biotic communities, of climate and landscape on individual ecosystem functions, for Sweden (model 1 to 3) and Madagascar (model 4 to 6).

| **Model** | **Response** | **Predictors** |
| --- | --- | --- |
| **1:** With species richness as a biotic driver, Sweden | Herbivory | Temperature + soil moisture + vegetation cover + species richness |
|  | Seed dispersal | Temperature + soil moisture + vegetation cover + species richness |
|  | Predation | Temperature + soil moisture + vegetation cover + species richness |
|  | Microbial decomposition | Temperature + soil moisture + leaf litter + species richness |
|  | Pollination | Temperature + soil moisture + vegetation cover + species richness |
|  | Species richness | Temperature + soil moisture + vegetation cover + leaf litter |
|  | Soil moisture | Temperature + vegetation cover + leaf litter |
|  | Leaf litter | Vegetation cover |
| **2:** With insect biomass as a biotic driver, Sweden | Herbivory | Temperature + soil moisture + vegetation cover + insect biomass |
|  | Seed dispersal | Temperature + soil moisture + vegetation cover + insect biomass |
|  | Predation | Temperature + soil moisture + vegetation cover + insect biomass |
|  | Microbial decomposition | Temperature + soil moisture + leaf litter + insect biomass |
|  | Pollination | Temperature + soil moisture + vegetation cover + insect biomass |
|  | Insect biomass | Temperature + soil moisture + vegetation cover + leaf litter |
|  | Soil moisture | Temperature + vegetation cover + leaf litter |
|  | Leaf litter | Vegetation cover |
| **3:** With community composition as a biotic driver, Sweden | Herbivory | Temperature + soil moisture + vegetation cover + community composition |
|  | Seed dispersal | Temperature + soil moisture + vegetation cover + community composition |
|  | Predation | Temperature + soil moisture + vegetation cover + community composition |
|  | Microbial decomposition | Temperature + soil moisture + leaf litter + community composition |
|  | Pollination | Temperature + soil moisture + vegetation cover + community composition |
|  | Community composition | Temperature + soil moisture + vegetation cover + leaf litter |
|  | Soil moisture | Temperature + vegetation cover + leaf litter |
|  | Leaf litter | Vegetation cover |
| **4: With species richness as a biotic driver, Madagascar** | Herbivory | Temperature + soil moisture + vegetation cover + species richness |
|  | Seed dispersal | Temperature + soil moisture + vegetation cover + species richness |
|  | Predation | Temperature + soil moisture + vegetation cover + species richness |
|  | Microbial decomposition | Temperature + soil moisture + leaf litter + species richness |
|  | Decomposition | Temperature + soil moisture + leaf litter + species richness |
|  | Species richness | Temperature + soil moisture + vegetation cover + leaf litter |
|  | Soil moisture | Temperature + vegetation cover + leaf litter |
|  | Leaf litter | Vegetation cover |
| **5: With insect biomass as a biotic driver, Madagascar** | Herbivory | Temperature + soil moisture + vegetation cover + insect biomass |
|  | Seed dispersal | Temperature + soil moisture + vegetation cover + insect biomass |
|  | Predation | Temperature + soil moisture + vegetation cover + insect biomass |
|  | Microbial decomposition | Temperature + soil moisture + leaf litter + insect biomass |
|  | Decomposition | Temperature + soil moisture + leaf litter + insect biomass |
|  | Insect biomass | Temperature + soil moisture + vegetation cover + leaf litter |
|  | Soil moisture | Temperature + vegetation cover + leaf litter |
|  | Leaf litter | Vegetation cover |
| **6: With community composition as a biotic driver, Madagascar** | Herbivory | Temperature + soil moisture + vegetation cover + community composition |
|  | Seed dispersal | Temperature + soil moisture + vegetation cover + community composition |
|  | Predation | Temperature + soil moisture + vegetation cover + community composition |
|  | Microbial decomposition | Temperature + soil moisture + leaf litter + community composition |
|  | Decomposition | Temperature + soil moisture + leaf litter + community composition |
|  | Community composition | Temperature + soil moisture + vegetation cover + leaf litter |
|  | Soil moisture | Temperature + vegetation cover + leaf litter |
|  | Leaf litter | Vegetation cover |

**Table S4.** Overview of fitted structural equation models to analyze the direct and indirect effects, i.e., via biotic communities, of climate and landscape on ecosystem multifunctionality. Fitted models were similar for Sweden and Madagascar.

| **Model** | **Response** | **Predictors** |
| --- | --- | --- |
| **1:** With species richness as biotic driver | Ecosystem multifunctionality | Temperature + soil moisture + vegetation cover + leaf litter + species richness |
|  | Species richness | Temperature + soil moisture + vegetation cover + leaf litter |
|  | Soil moisture | Temperature + vegetation cover + leaf litter |
|  | Leaf litter | Vegetation cover |
| **2:** With insect biomass as biotic driver | Ecosystem multifunctionality | Temperature + soil moisture + vegetation cover + leaf litter + insect biomass |
|  | Insect biomass | Temperature + soil moisture + vegetation cover + leaf litter |
|  | Soil moisture | Temperature + vegetation cover + leaf litter |
|  | Leaf litter | Vegetation cover |
| **3:** With community composition as biotic driver | Ecosystem multifunctionality | Temperature + soil moisture + vegetation cover + leaf litter + community composition |
|  | Community composition | Temperature + soil moisture + vegetation cover + leaf litter |
|  | Soil moisture | Temperature + vegetation cover + leaf litter |
|  | Leaf litter | Vegetation cover |

**Table S5.**
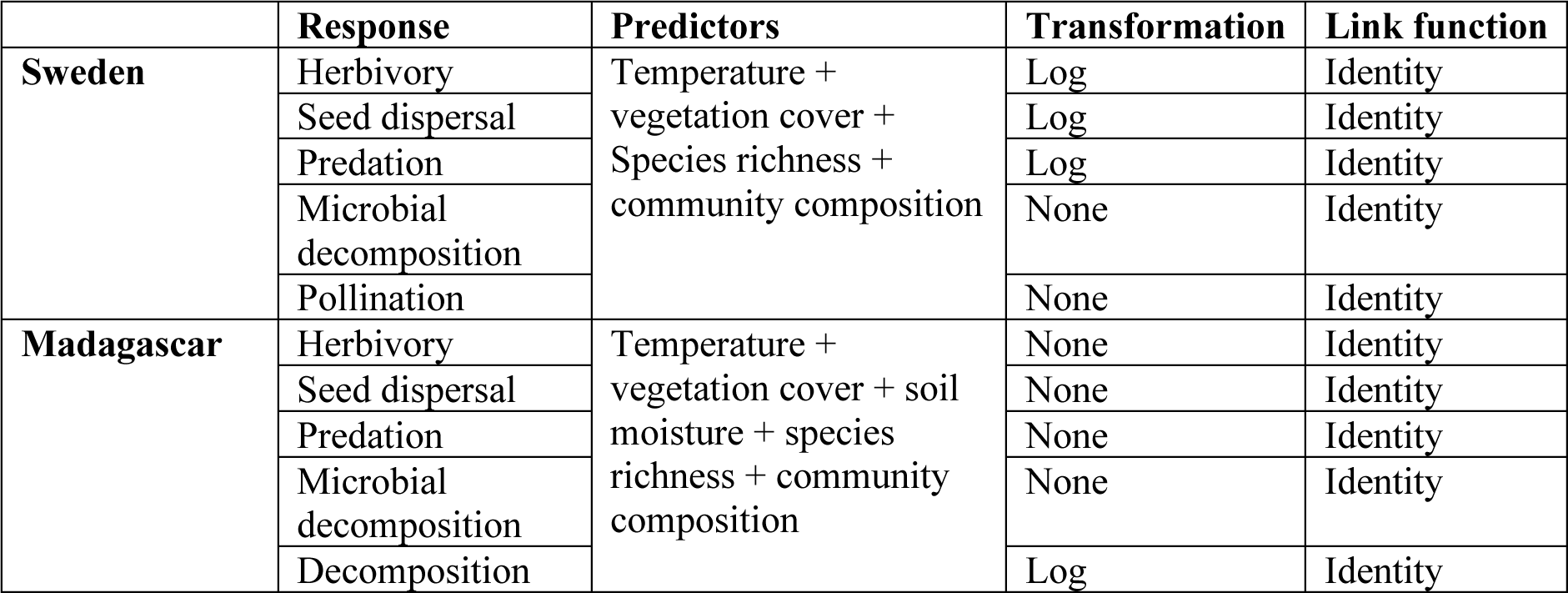
Overview of fitted models on the effect of climate, landscape and the biotic community on individual ecosystem functions, from which we obtained the residuals, for Sweden and Madagascar.

|  | **Response** | **Predictors** | **Transformation** | **Link function** |
| --- | --- | --- | --- | --- |
| **Sweden** | Herbivory | Temperature +<br>vegetation cover +<br>Species richness +<br>community composition | Log | Identity |
|  | Seed dispersal |  | Log | Identity |
|  | Predation |  | Log | Identity |
|  | Microbial decomposition |  | None | Identity |
|  | Pollination |  | None | Identity |
| **Madagascar** | Herbivory | Temperature +<br>vegetation cover + soil<br>moisture + species<br>richness + community<br>composition | None | Identity |
|  | Seed dispersal |  | None | Identity |
|  | Predation |  | None | Identity |
|  | Microbial decomposition |  | None | Identity |
|  | Decomposition |  | Log | Identity |

**Table S6.**
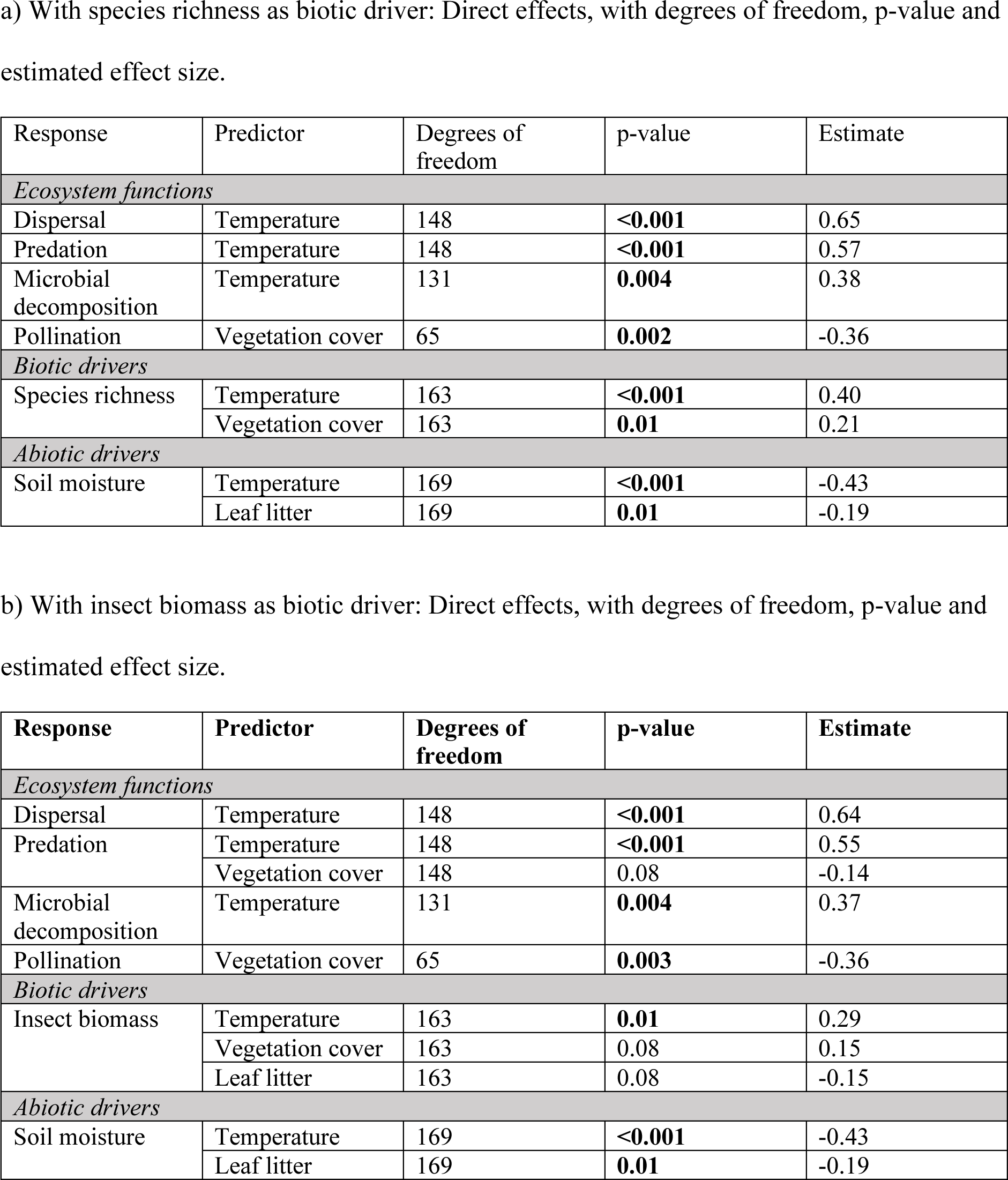
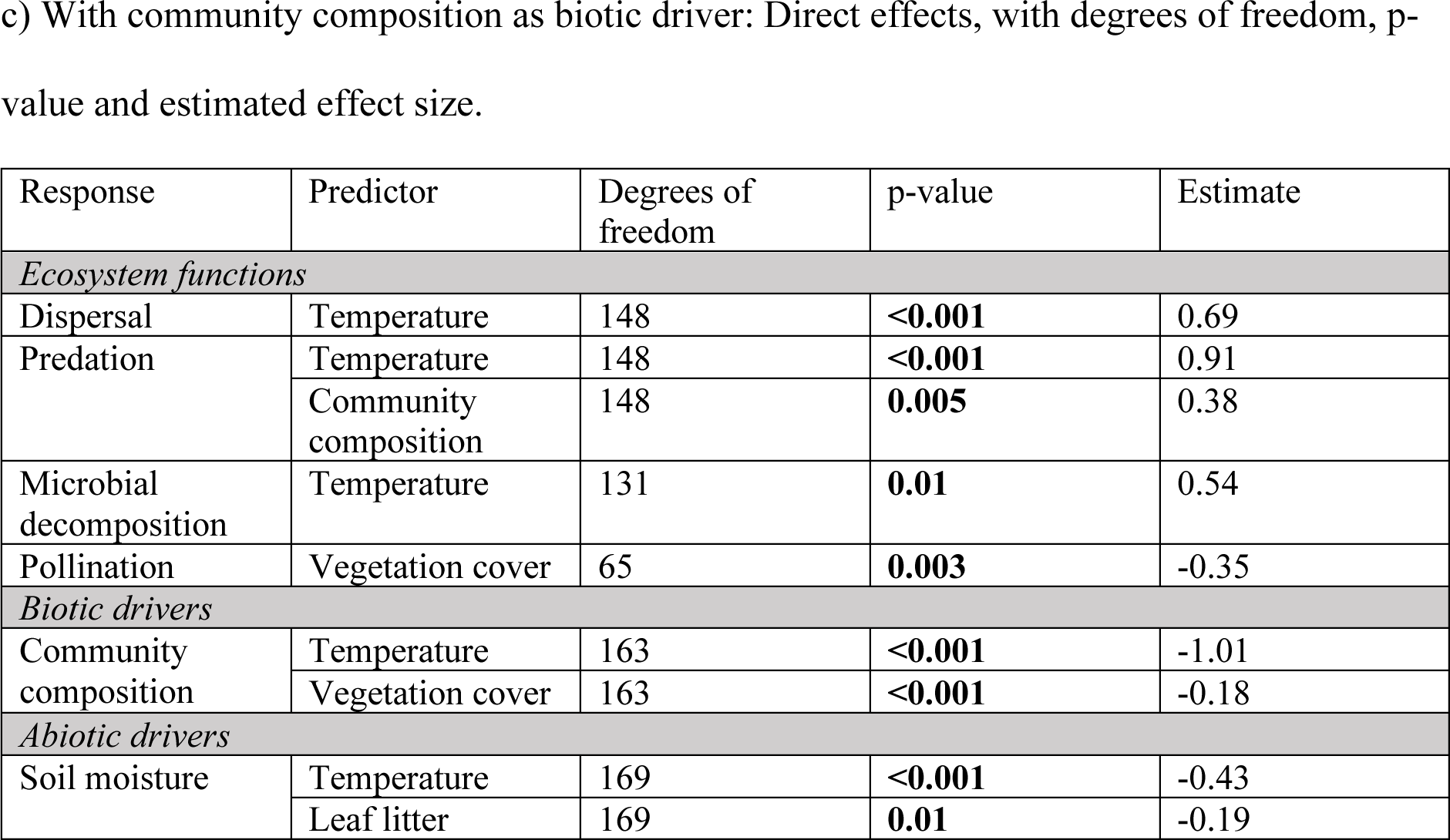
Statistical output from SEM models on individual ecosystem functions in Sweden. Shown are significant (p < 0.05) and near significant (0.05 < p < 0.1) effects.

| Response | Predictor | Degrees of freedom | p-value | Estimate |
| --- | --- | --- | --- | --- |
| *Ecosystem functions* |  |  |  |  |
| Dispersal | Temperature | 148 | **<0.001** | 0.65 |
| Predation | Temperature | 148 | **<0.001** | 0.57 |
| Microbial decomposition | Temperature | 131 | **0.004** | 0.38 |
| Pollination | Vegetation cover | 65 | **0.002** | -0.36 |
| *Biotic drivers* |  |  |  |  |
| Species richness | Temperature | 163 | **<0.001** | 0.40 |
|  | Vegetation cover | 163 | **0.01** | 0.21 |
| *Abiotic drivers* |  |  |  |  |
| Soil moisture | Temperature | 169 | **<0.001** | -0.43 |
|  | Leaf litter | 169 | **0.01** | -0.19 |

b) With insect biomass as biotic driver: Direct effects, with degrees of freedom, p-value and estimated effect size.
| Response | Predictor | Degrees of freedom | p-value | Estimate |
| --- | --- | --- | --- | --- |
| *Ecosystem functions* |  |  |  |  |
| Dispersal | Temperature | 148 | **<0.001** | 0.64 |
| Predation | Temperature | 148 | **<0.001** | 0.55 |
|  | Vegetation cover | 148 | 0.08 | -0.14 |
| Microbial decomposition | Temperature | 131 | **0.004** | 0.37 |
| Pollination | Vegetation cover | 65 | **0.003** | -0.36 |
| *Biotic drivers* |  |  |  |  |
| Insect biomass | Temperature | 163 | **0.01** | 0.29 |
|  | Vegetation cover | 163 | 0.08 | 0.15 |
|  | Leaf litter | 163 | 0.08 | -0.15 |
| *Abiotic drivers* |  |  |  |  |
| Soil moisture | Temperature | 169 | **<0.001** | -0.43 |
|  | Leaf litter | 169 | **0.01** | -0.19 |
| *Ecosystem functions* |  |  |  |  |
| Dispersal | Temperature | 148 | **<0.001** | 0.69 |
| Predation | Temperature | 148 | **<0.001** | 0.91 |
|  | Community composition | 148 | **0.005** | 0.38 |
| Microbial decomposition | Temperature | 131 | **0.01** | 0.54 |
| Pollination | Vegetation cover | 65 | **0.003** | -0.35 |
| *Biotic drivers* |  |  |  |  |
| Community composition | Temperature | 163 | **<0.001** | -1.01 |
|  | Vegetation cover | 163 | **<0.001** | -0.18 |
| *Abiotic drivers* |  |  |  |  |
| Soil moisture | Temperature | 169 | **<0.001** | -0.43 |
|  | Leaf litter | 169 | **0.01** | -0.19 |

**Table S7.**
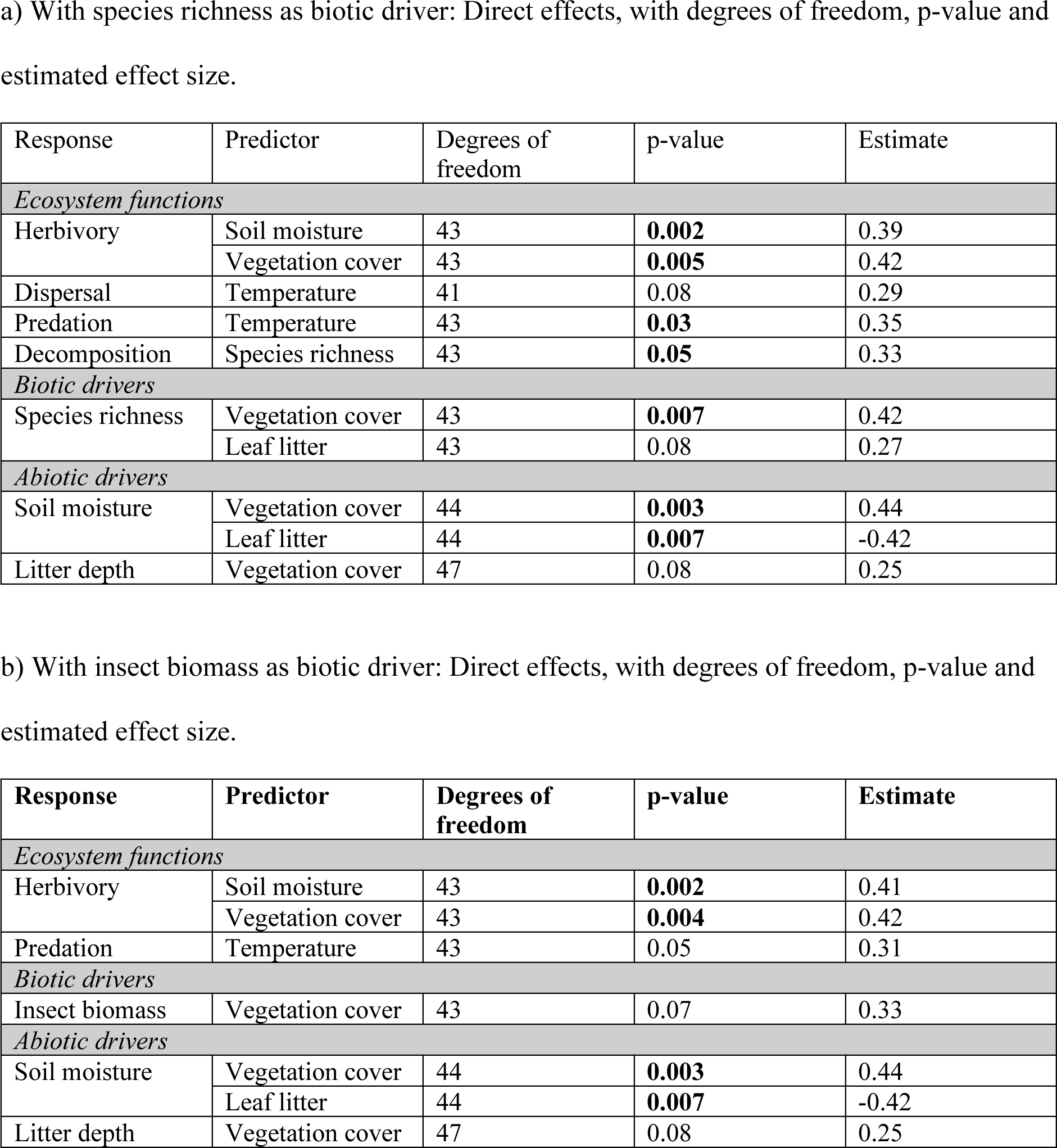
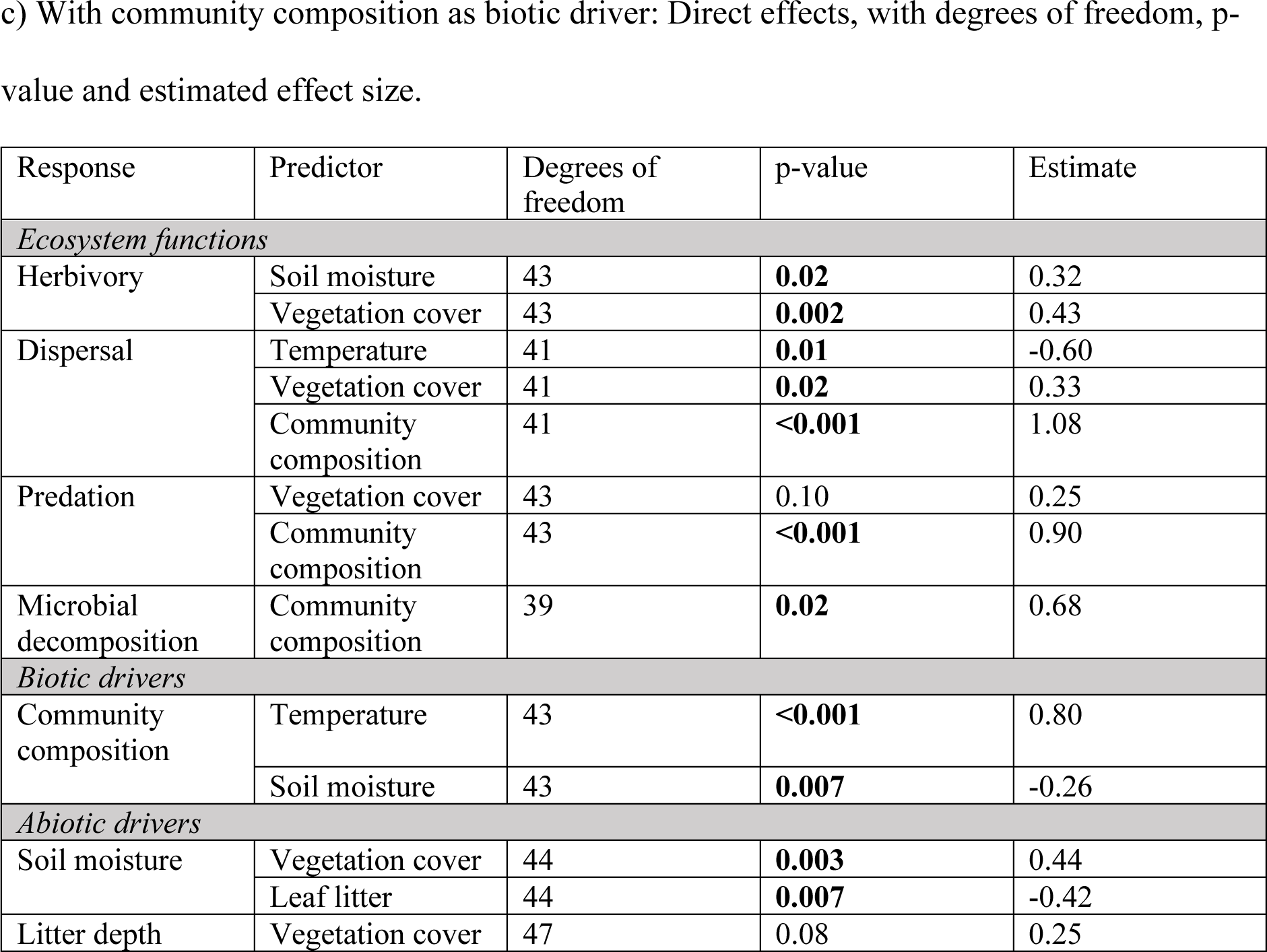
Statistical output from SEM models on individual ecosystem functions in Madagascar. Shown are significant (p < 0.05) and near significant (0.05 < p < 0.1) effects.

| Response | Predictor | Degrees of freedom | p-value | Estimate |
| --- | --- | --- | --- | --- |
| *Ecosystem functions* |  |  |  |  |
| Herbivory | Soil moisture | 43 | **0.002** | 0.39 |
|  | Vegetation cover | 43 | **0.005** | 0.42 |
| Dispersal | Temperature | 41 | 0.08 | 0.29 |
| Predation | Temperature | 43 | **0.03** | 0.35 |
| Decomposition | Species richness | 43 | **0.05** | 0.33 |
| *Biotic drivers* |  |  |  |  |
| Species richness | Vegetation cover | 43 | **0.007** | 0.42 |
|  | Leaf litter | 43 | 0.08 | 0.27 |
| *Abiotic drivers* |  |  |  |  |
| Soil moisture | Vegetation cover | 44 | **0.003** | 0.44 |
|  | Leaf litter | 44 | **0.007** | -0.42 |
| Litter depth | Vegetation cover | 47 | 0.08 | 0.25 |

b) With insect biomass as biotic driver: Direct effects, with degrees of freedom, p-value and estimated effect size.
| Response | Predictor | Degrees of freedom | p-value | Estimate |
| --- | --- | --- | --- | --- |
| *Ecosystem functions* |  |  |  |  |
| Herbivory | Soil moisture | 43 | **0.002** | 0.41 |
|  | Vegetation cover | 43 | **0.004** | 0.42 |
| Predation | Temperature | 43 | 0.05 | 0.31 |
| *Biotic drivers* |  |  |  |  |
| Insect biomass | Vegetation cover | 43 | 0.07 | 0.33 |
| *Abiotic drivers* |  |  |  |  |
| Soil moisture | Vegetation cover | 44 | **0.003** | 0.44 |
|  | Leaf litter | 44 | **0.007** | -0.42 |
| Litter depth | Vegetation cover | 47 | 0.08 | 0.25 |
| *Ecosystem functions* |  |  |  |  |
| Herbivory | Soil moisture | 43 | **0.02** | 0.32 |
|  | Vegetation cover | 43 | **0.002** | 0.43 |
| Dispersal | Temperature | 41 | **0.01** | -0.60 |
|  | Vegetation cover | 41 | **0.02** | 0.33 |
|  | Community composition | 41 | **<0.001** | 1.08 |
| Predation | Vegetation cover | 43 | 0.10 | 0.25 |
|  | Community composition | 43 | **<0.001** | 0.90 |
| Microbial decomposition | Community composition | 39 | **0.02** | 0.68 |
| *Biotic drivers* |  |  |  |  |
| Community composition | Temperature | 43 | **<0.001** | 0.80 |
|  | Soil moisture | 43 | **0.007** | -0.26 |
| *Abiotic drivers* |  |  |  |  |
| Soil moisture | Vegetation cover | 44 | **0.003** | 0.44 |
|  | Leaf litter | 44 | **0.007** | -0.42 |
| Litter depth | Vegetation cover | 47 | 0.08 | 0.25 |

**Table S8.**
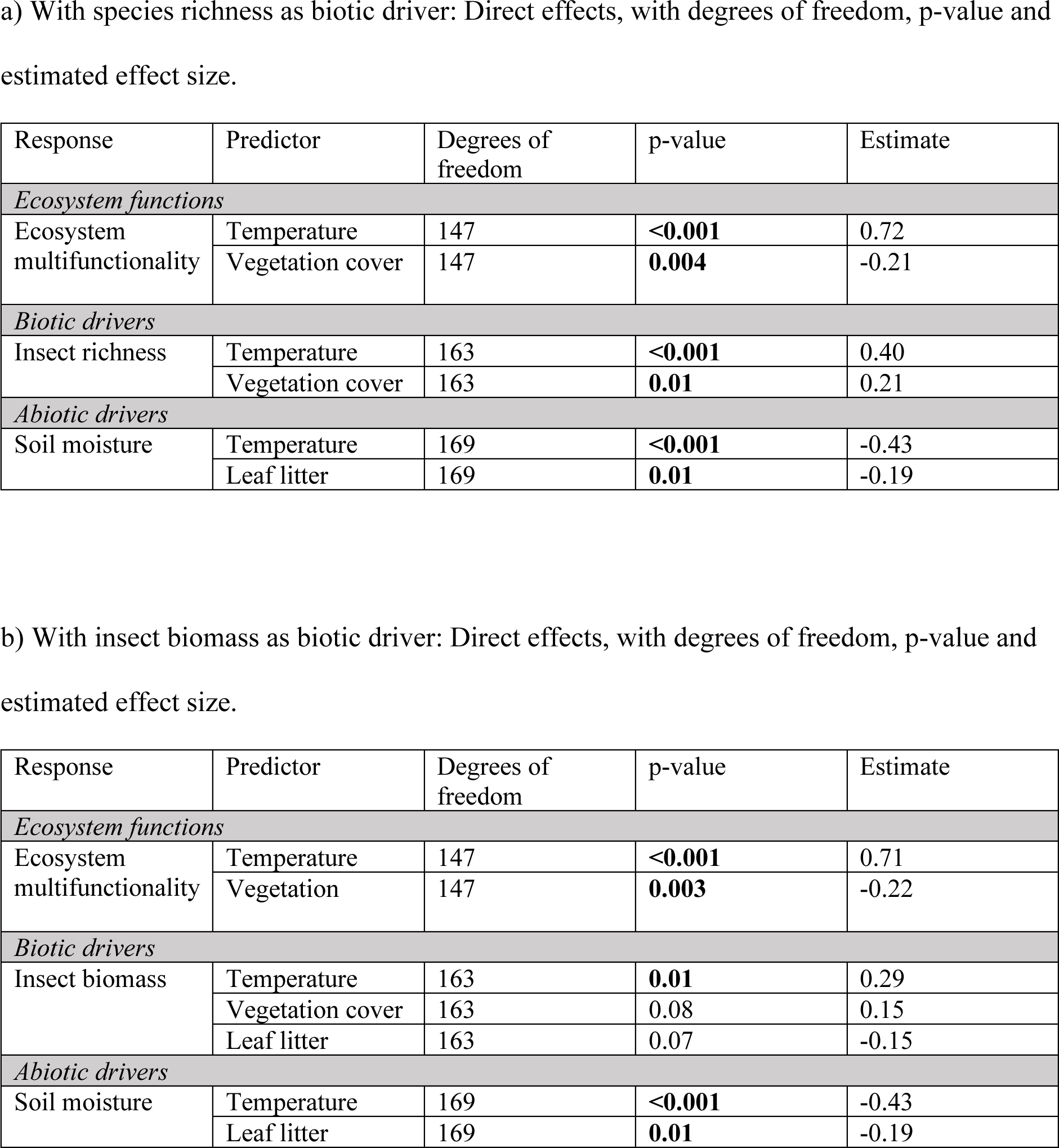
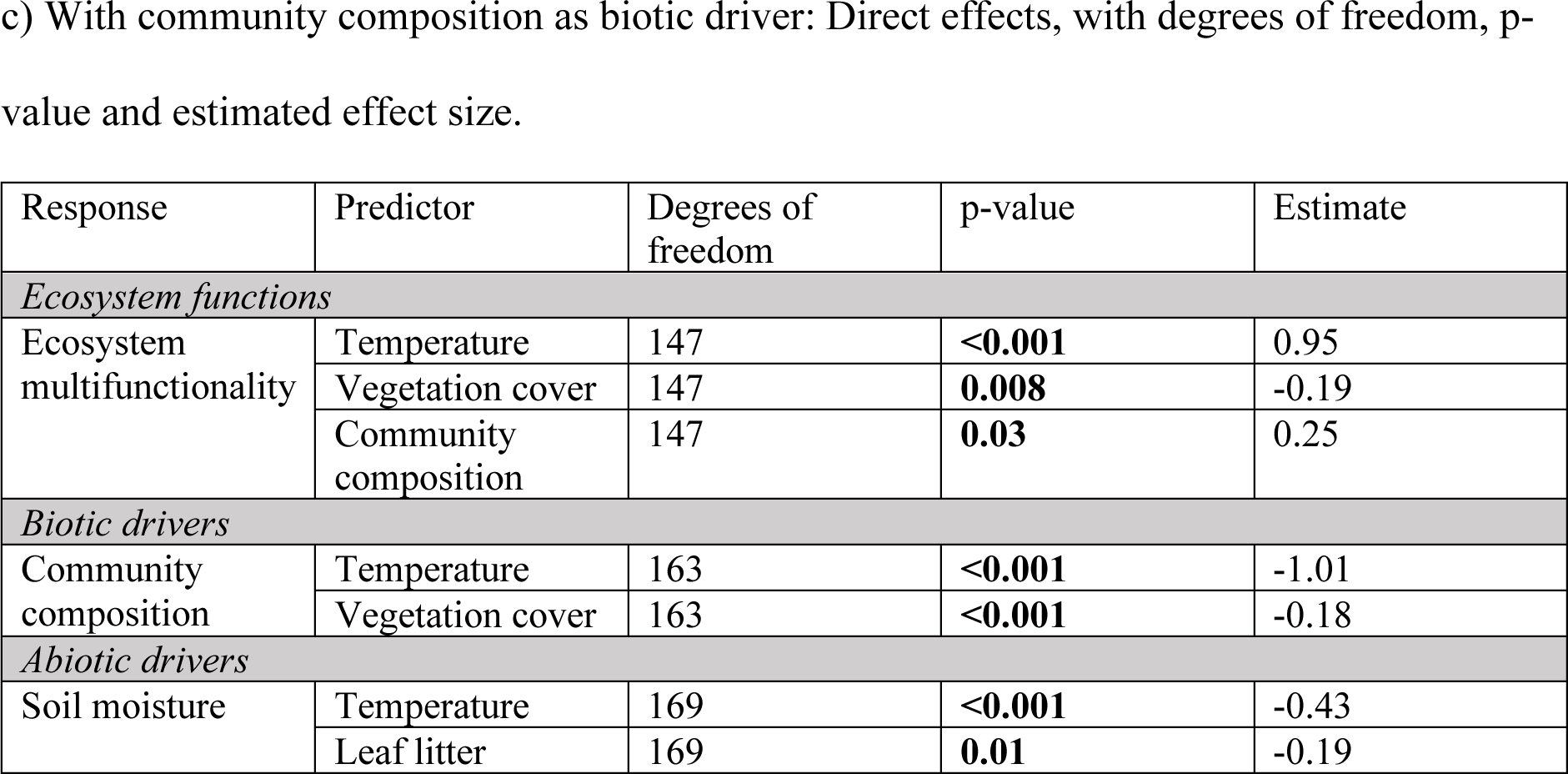
Statistical output from SEM models on ecosystem multifunctionality in Sweden. Shown are significant (p < 0.05) and near significant (0.05 < p < 0.1) effects.

| Response | Predictor | Degrees of freedom | p-value | Estimate |
| --- | --- | --- | --- | --- |
| *Ecosystem functions* |  |  |  |  |
| Ecosystem multifunctionality | Temperature | 147 | **<0.001** | 0.72 |
|  | Vegetation cover | 147 | **0.004** | -0.21 |
| *Biotic drivers* |  |  |  |  |
| Insect richness | Temperature | 163 | **<0.001** | 0.40 |
|  | Vegetation cover | 163 | **0.01** | 0.21 |
| *Abiotic drivers* |  |  |  |  |
| Soil moisture | Temperature | 169 | **<0.001** | -0.43 |
|  | Leaf litter | 169 | **0.01** | -0.19 |

b) With insect biomass as biotic driver: Direct effects, with degrees of freedom, p-value and estimated effect size.
| Response | Predictor | Degrees of freedom | p-value | Estimate |
| --- | --- | --- | --- | --- |
| *Ecosystem functions* |  |  |  |  |
| Ecosystem multifunctionality | Temperature | 147 | **<0.001** | 0.71 |
|  | Vegetation | 147 | **0.003** | -0.22 |
| *Biotic drivers* |  |  |  |  |
| Insect biomass | Temperature | 163 | **0.01** | 0.29 |
|  | Vegetation cover | 163 | 0.08 | 0.15 |
|  | Leaf litter | 163 | 0.07 | -0.15 |
| *Abiotic drivers* |  |  |  |  |
| Soil moisture | Temperature | 169 | **<0.001** | -0.43 |
|  | Leaf litter | 169 | **0.01** | -0.19 |
| *Ecosystem functions* |  |  |  |  |
| Ecosystem multifunctionality | Temperature | 147 | **<0.001** | 0.95 |
|  | Vegetation cover | 147 | **0.008** | -0.19 |
|  | Community composition | 147 | **0.03** | 0.25 |
| *Biotic drivers* |  |  |  |  |
| Community composition | Temperature | 163 | **<0.001** | -1.01 |
|  | Vegetation cover | 163 | **<0.001** | -0.18 |
| *Abiotic drivers* |  |  |  |  |
| Soil moisture | Temperature | 169 | **<0.001** | -0.43 |
|  | Leaf litter | 169 | **0.01** | -0.19 |

**Table S9.**
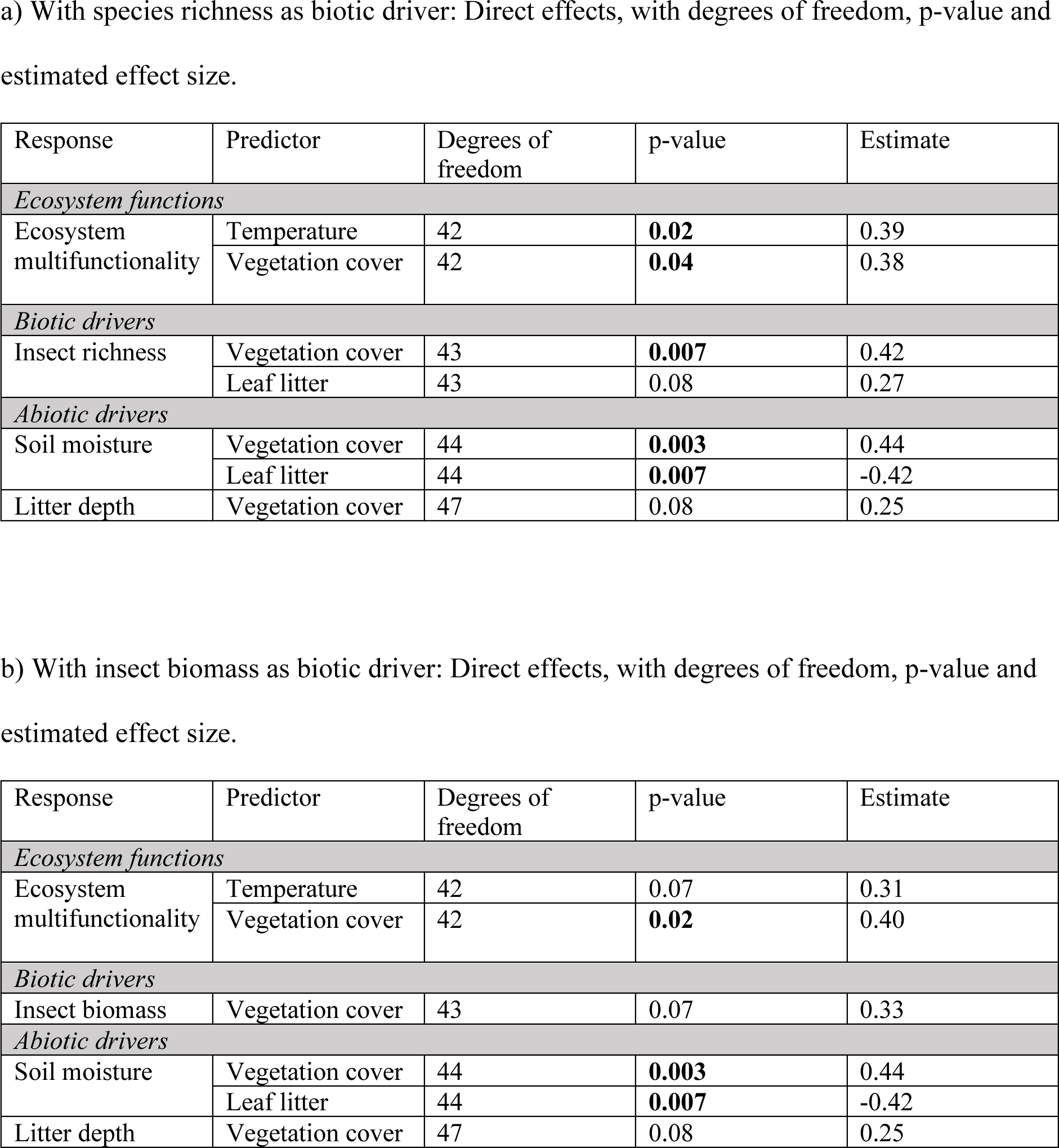
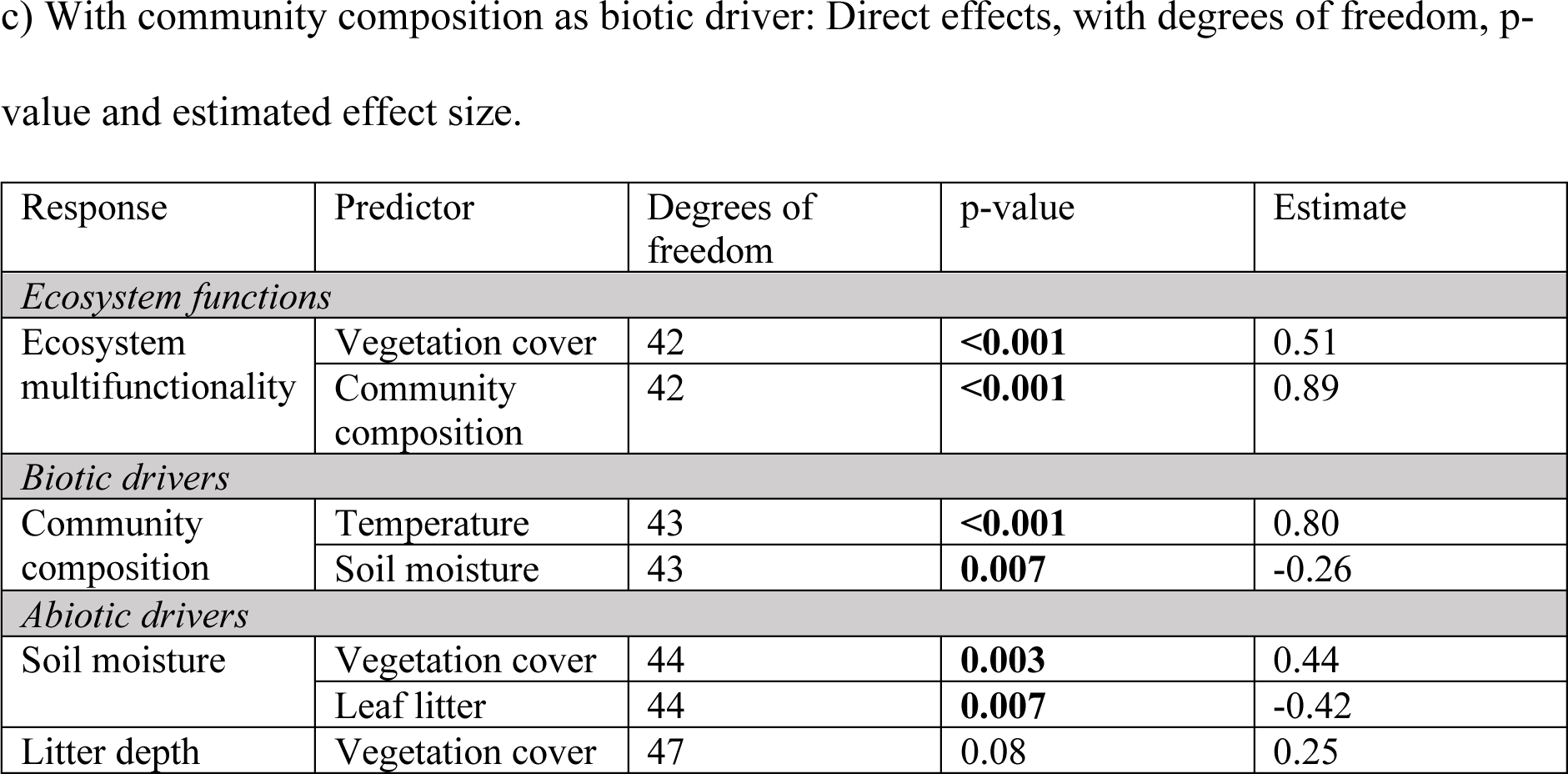
Statistical output from SEM models on ecosystem multifunctionality in Madagascar. Shown are significant (p < 0.05) and near significant (0.05 < p < 0.1) effects.

| Response | Predictor | Degrees of freedom | p-value | Estimate |
| --- | --- | --- | --- | --- |
| *Ecosystem functions* |  |  |  |  |
| Ecosystem multifunctionality | Temperature | 42 | **0.02** | 0.39 |
|  | Vegetation cover | 42 | **0.04** | 0.38 |
| *Biotic drivers* |  |  |  |  |
| Insect richness | Vegetation cover | 43 | **0.007** | 0.42 |
|  | Leaf litter | 43 | 0.08 | 0.27 |
| *Abiotic drivers* |  |  |  |  |
| Soil moisture | Vegetation cover | 44 | **0.003** | 0.44 |
|  | Leaf litter | 44 | **0.007** | -0.42 |
| Litter depth | Vegetation cover | 47 | 0.08 | 0.25 |

b) With insect biomass as biotic driver: Direct effects, with degrees of freedom, p-value and estimated effect size.
| Response | Predictor | Degrees of freedom | p-value | Estimate |
| --- | --- | --- | --- | --- |
| *Ecosystem functions* |  |  |  |  |
| Ecosystem multifunctionality | Temperature | 42 | 0.07 | 0.31 |
|  | Vegetation cover | 42 | **0.02** | 0.40 |
| *Biotic drivers* |  |  |  |  |
| Insect biomass | Vegetation cover | 43 | 0.07 | 0.33 |
| *Abiotic drivers* |  |  |  |  |
| Soil moisture | Vegetation cover | 44 | **0.003** | 0.44 |
|  | Leaf litter | 44 | **0.007** | -0.42 |
| Litter depth | Vegetation cover | 47 | 0.08 | 0.25 |
| *Ecosystem functions* |  |  |  |  |
| Ecosystem multifunctionality | Vegetation cover | 42 | **<0.001** | 0.51 |
|  | Community composition | 42 | **<0.001** | 0.89 |
| *Biotic drivers* |  |  |  |  |
| Community composition | Temperature | 43 | **<0.001** | 0.80 |
|  | Soil moisture | 43 | **0.007** | -0.26 |
| *Abiotic drivers* |  |  |  |  |
| Soil moisture | Vegetation cover | 44 | **0.003** | 0.44 |
|  | Leaf litter | 44 | **0.007** | -0.42 |
| Litter depth | Vegetation cover | 47 | 0.08 | 0.25 |

**Figure S1.**
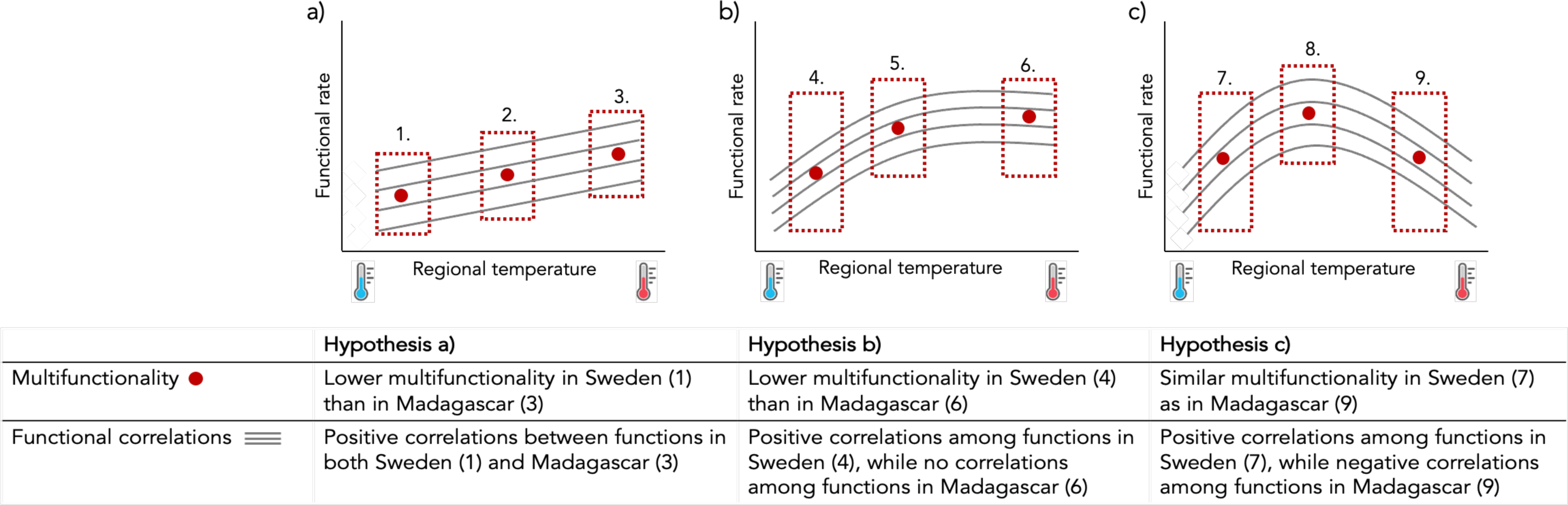
Hypotheses on the effect of regional temperature on functional rates of several ecosystem functions. Panels a)-c) illustrate different hypothesized patterns based on regional temperatures, and the table below describes how patterns in Sweden and Madagascar will be observed under these different scenarios. In panel a), regional temperature has a positive linear effect on functional rates. When looking at local patterns, all sites will show a positive correlation among functions, and colder sites (1) will have lower multifunctionality compared to warmer sites (2, 3). In panel b), regional temperature positively affects functional rates, but the positive effect of temperature turns neutral towards the warmest regions. Hence, correlations among functions will look different depending on the sampled location, with positive correlations among functions in the coldest locations (4), weaker positive correlations in locations with intermediate temperature (5), and no correlations among functions in the warmest locations (6). In this scenario, multifunctionality will also increase from the coldest (4) to the warmest locations (6). In panel c), regional temperature positively affects functional rates up to a certain threshold, after which temperature has a negative effect on functional rates. In this scenario, positive correlations among functions will be observed in colder locations (7), no correlations will be observed in locations with intermediate temperature (8), and negative correlations will be observed in locations with the warmest regional temperatures (9). Here, ecosystem multifunctionality will be highest in locations with intermediate temperatures (8), and lowest in locations with colder or warmer temperatures (7, 9).

**Figure S2.**
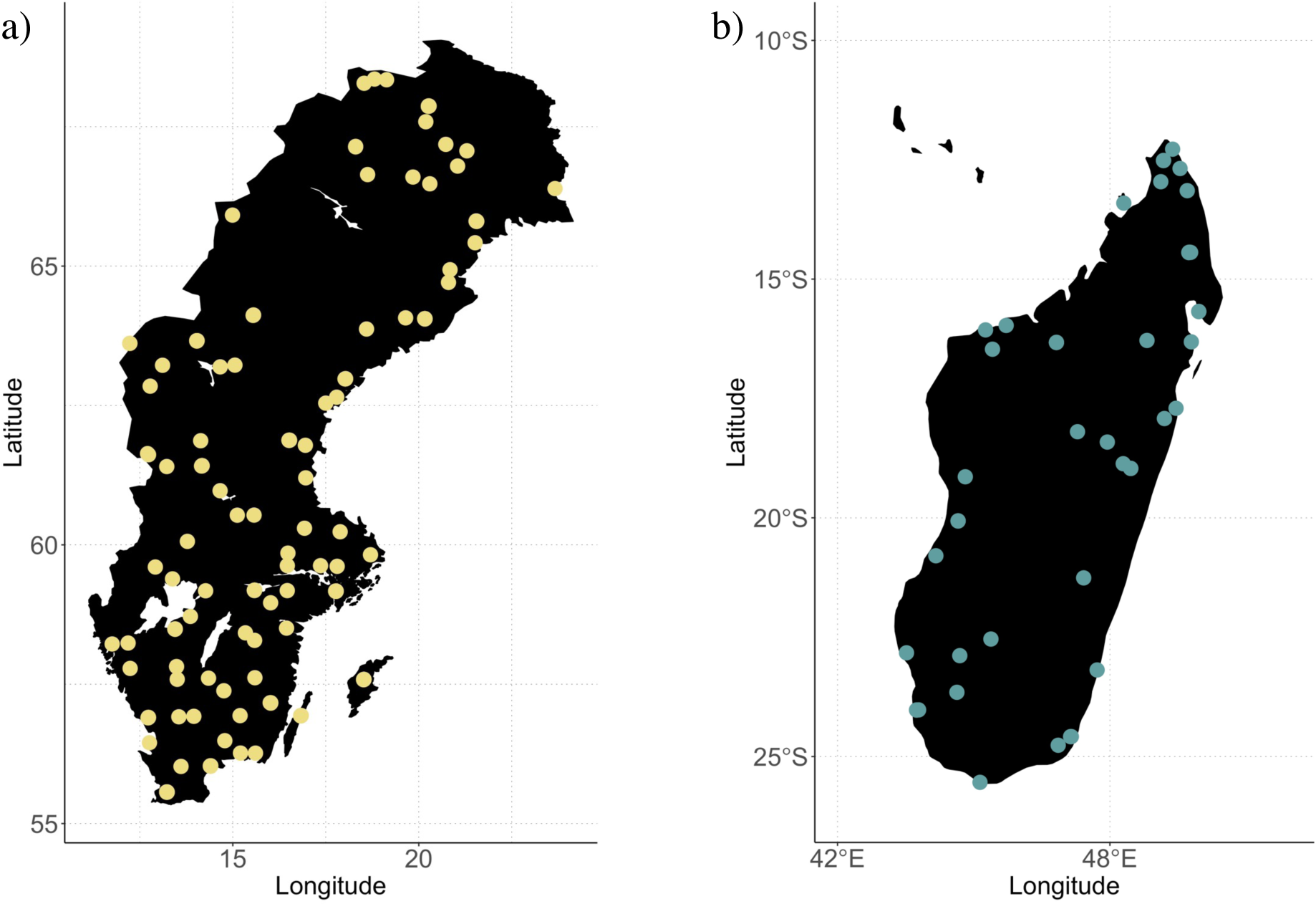
Map with locations at which ecosystem functions were measured, in (a) Sweden and (b) Madagascar.

**Figure S3.**
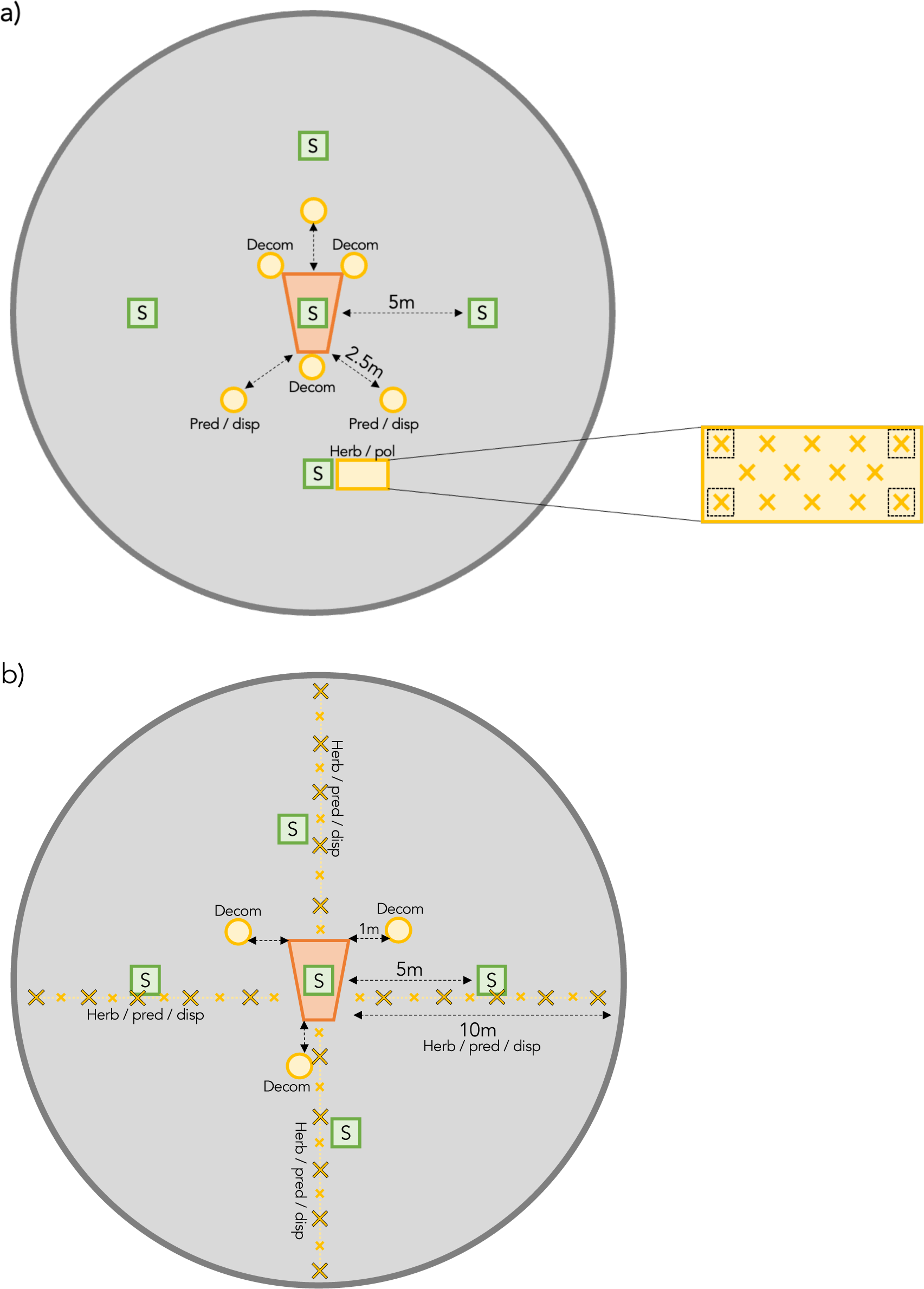
Experimental setup for measurements of ecosystem functions, climate and the insect community in **a)** Sweden and **b)** Madagascar. The experimental area is indicated as a grey circle (r = 10 m). In the center of the experimental area, a Malaise trap was placed to collect insects (orange). Experiments on ecosystem functions are indicated in yellow, and for **a)** Sweden, include: i) three locations with dispersal and predation experiments (“pred / disp”, circle) at 2.5 meters from the trap, ii) 3 locations with microbial decomposition experiments (“decom”, circle) directly at the corners of the trap, and iii) 1 spot with pollination and herbivory experiments (“herb / pol”, rectangle) at ∼5 meters from the trap. For the herbivory and pollination experiments, strawberry plants were placed in the corners of a 15 L soil bag (crosses surrounded by dashed black square), and willow sticks were placed in rows in the center of the soil bag (crosses without square). In **b)** Madagascar, ecosystem function experiments included: i) four 10 m transects with dispersal and predation experiments every 2m (“pred / disp”, big crosses), ii) four 10 m transects with leaf collection every 1 m for herbivory assessments (“herb”, big crosses and small crosses), iii) 3 locations with microbial decomposition experiments (“decom”, circle) with lamina baits and tea bags, at 1 m distance from the corners of the trap. Measurements on local climate in **a)** and **b)** are indicated in green, and include measurements on soil moisture and leaf litter at five locations (squares, “S”) – one measurement at the Malaise trap, and four at five meters distance from the Malaise trap in different directions. For a summary of functions and methods measured per region, see Table S1.

**Figure S4.**
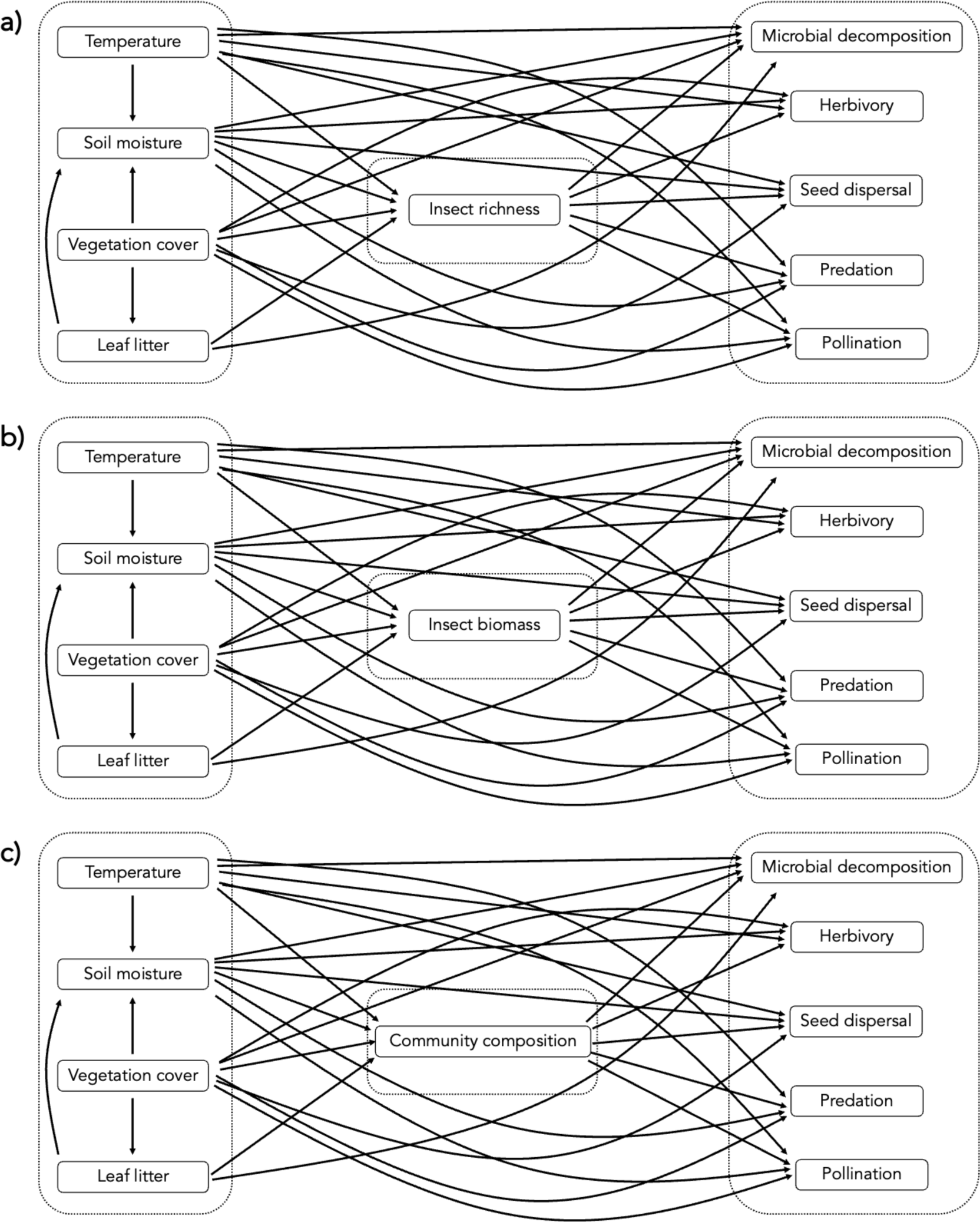
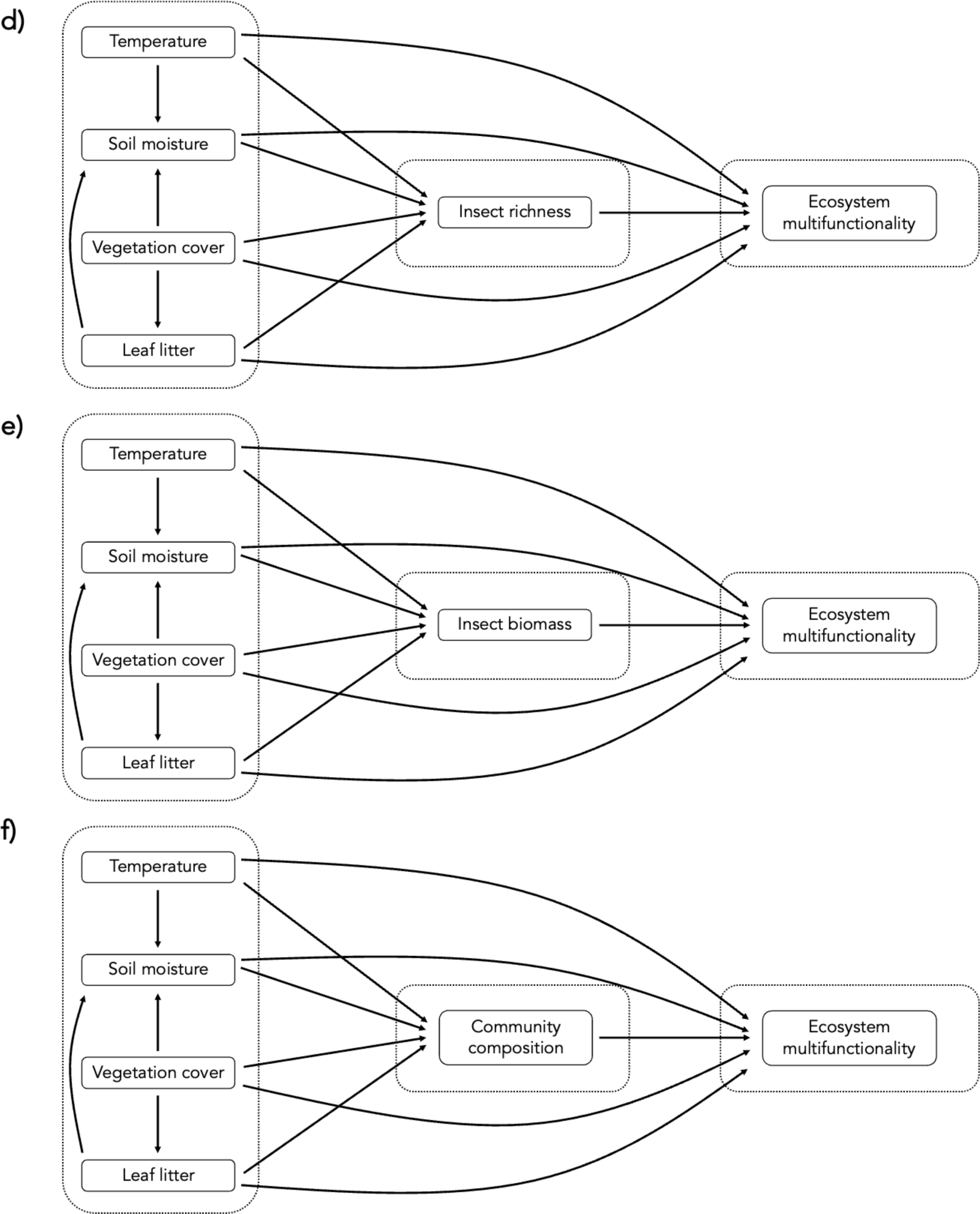
Conceptual models on the effect of climate, landscape and biotic communities on (a-c) individual ecosystem functions and (d-f) ecosystem multifunctionality in Sweden, with one model presented per panel (table S3). As biotic predictors, we used species richness for the models in panels a) and d), insect biomass for the models in panel b) and e), and community composition for the models in panels c) and f). Observed variables are presented in boxes, and directional relations as included in the SEM model are indicated with arrows.

**Figure S5.**
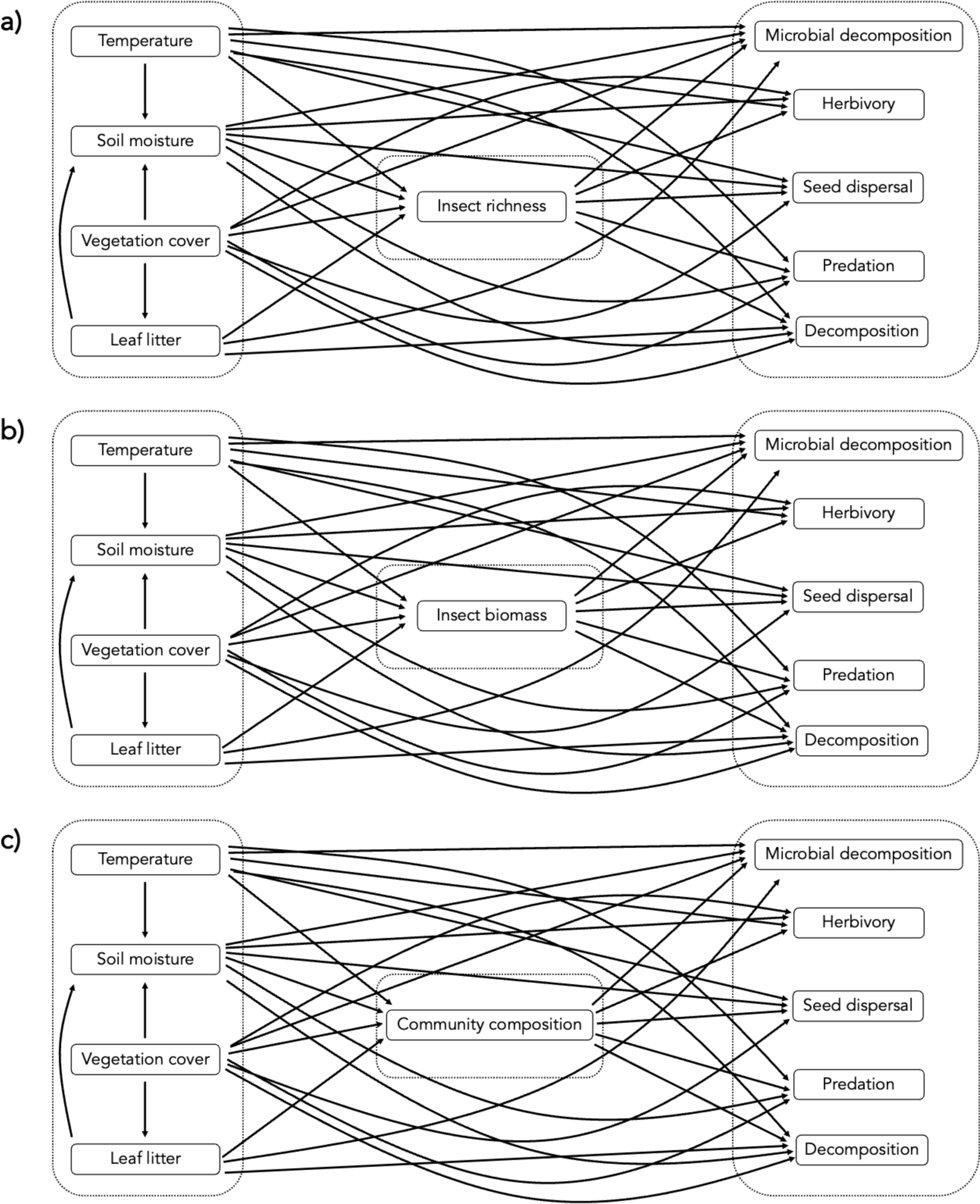
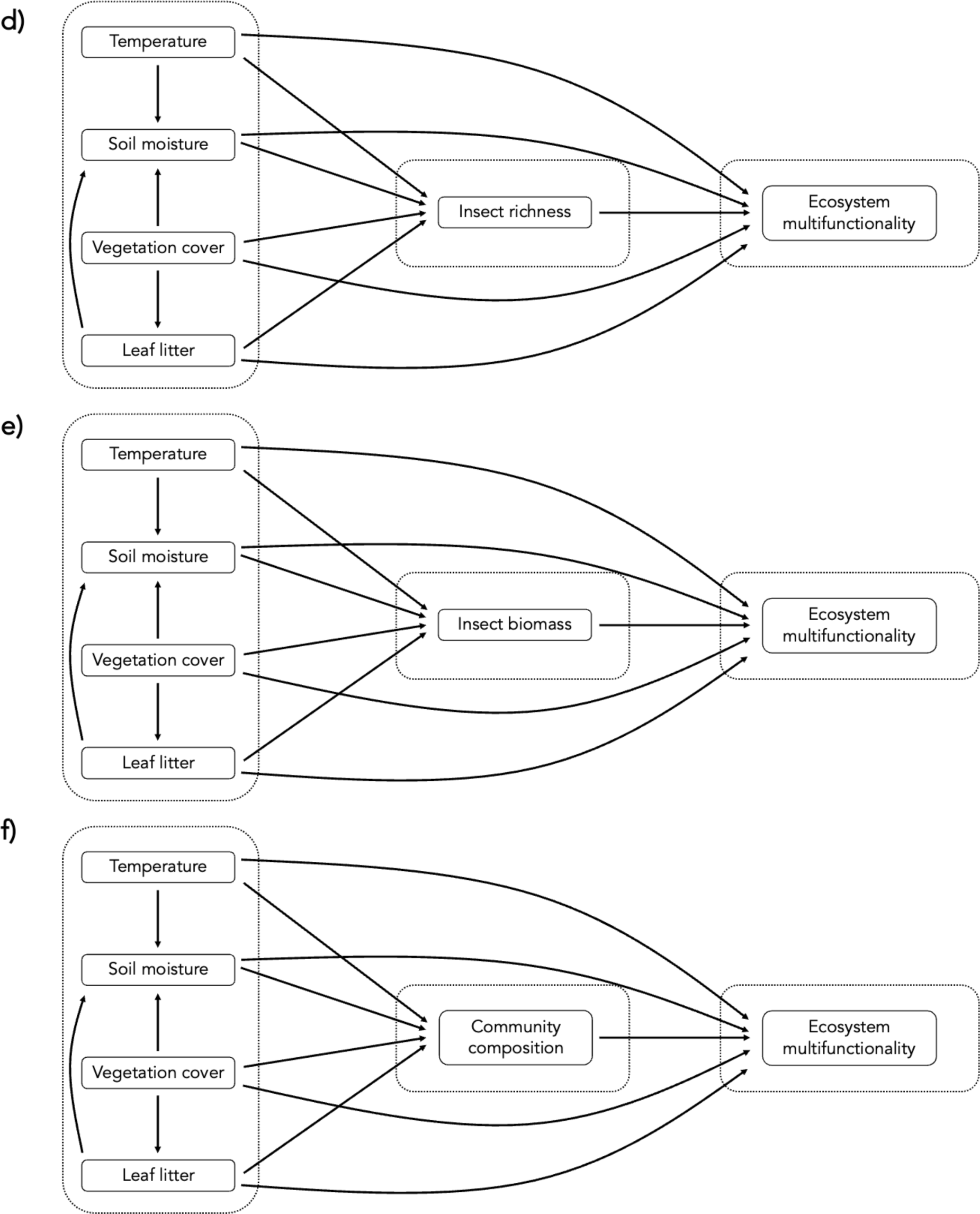
Conceptual models on the effect of climate, landscape and biotic communities on (a-c) individual ecosystem functions and (d-f) ecosystem multifunctionality in Madagascar, with one model presented per panel (table S3). As biotic predictors, we used species richness for the models in panels a) and d), insect biomass for the models in panel b) and e), and community composition for the models in panels c) and f). Observed variables are presented in boxes, and directional relations as included in the SEM model are indicated with arrows.

**Figure S6.**
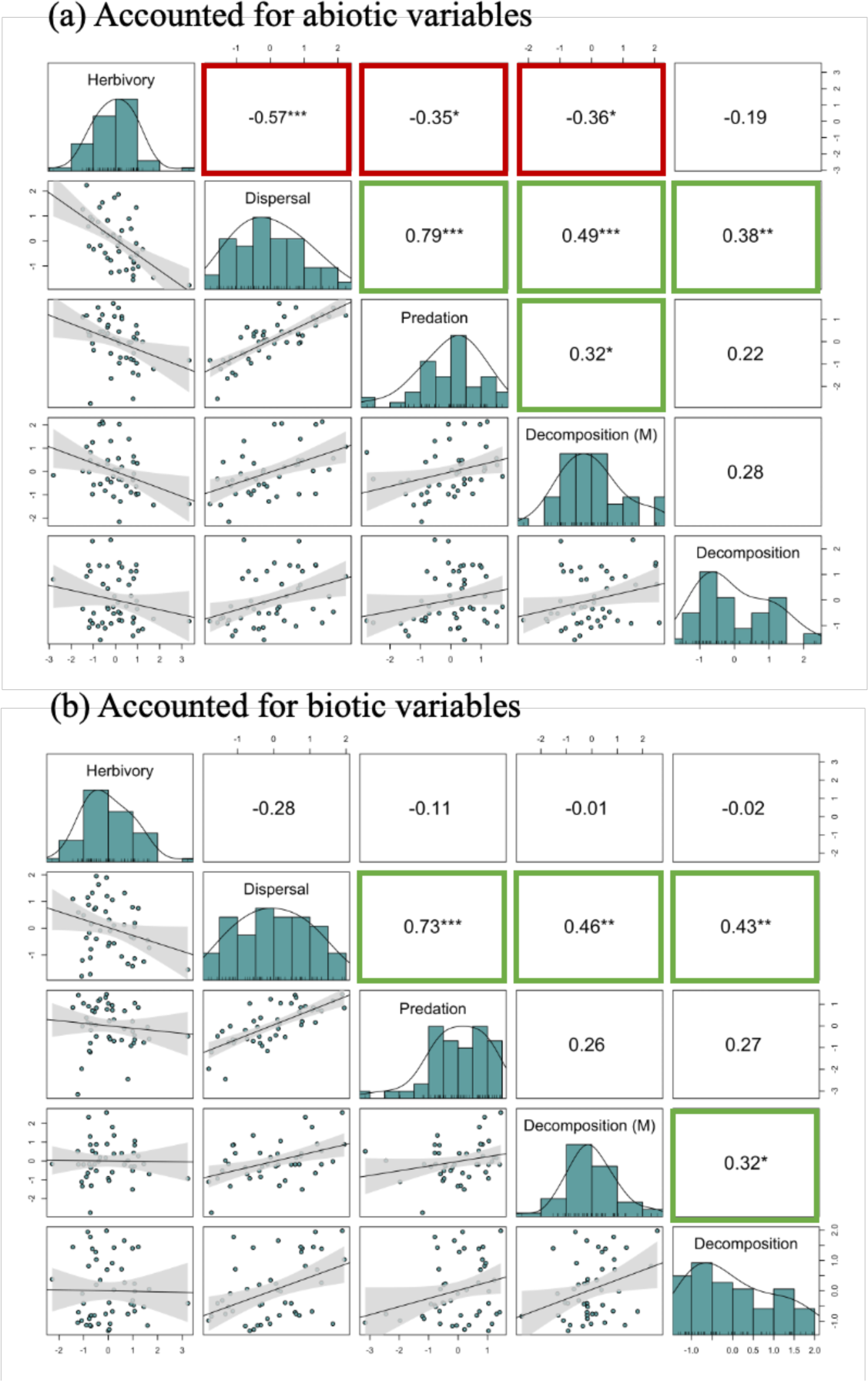
Residual correlations among ecosystem functions in Madagascar, after accounting for the main structuring (a) abiotic and (b) biotic forces identified by SEMs. Ecosystem functions include herbivory, seed dispersal, predation, microbial decomposition (“M”) and decomposition by invertebrates and microbes. Linear correlations between ecosystem functions are presented in the lower left corner of the panels, and the Pearson correlation coefficient (r) with stars of significance (* p < 0.05, ** p < 0.01, *** p < 0.001) in the upper right corner of the panels. Green-bordered squares indicate significantly positive correlations, and red-bordered squares indicate significantly negative correlations. The histograms in the diagonal present the distribution of the data for all ecosystem functions.

## Notes

### Competing Interest Statement

The authors have declared no competing interest.

